# Episodic rhythmicity is generated by a distributed neural network in the developing mammalian spinal cord

**DOI:** 10.1101/2024.07.26.605187

**Authors:** Jonathan J. Milla-Cruz, Adam P. Lognon, Michelle A. Tran, Stephanie A. Di Vito, Carlotta Löer, Anchita Shonak, Matthew J. Broadhead, Gareth B. Miles, Simon A. Sharples, Patrick J. Whelan

## Abstract

Spinal circuits produce diverse motor outputs that coordinate the rhythm and pattern of locomotor movements. Despite the episodic nature of these behaviours, the neural mechanisms encoding these episodes are not well understood. This study investigated mechanisms producing episodic rhythms evoked by dopamine in isolated neonatal mouse spinal cords. Dopamine-induced rhythms were primarily synchronous and propagated rostro-caudally across spinal segments, with occasional asynchronous episodes. Electrical stimulation of the L5 dorsal root could entrain episodes across segments, indicating afferent control of the rhythm generator and a distributed rostro-caudal network. Episodic activity was observed in isolated thoracic or sacral segments after full spinal transection or bilateral ventrolateral funiculus (VLF) lesions, suggesting a distributed network coupled via VLF projections. Rhythmicity was recorded from axons projecting through the VLF and dorsal roots, but not from cholinergic recurrent excitation via motoneurons or isolated dorsal inhibitory circuits. The data suggest episodic rhythmicity is generated by a flexibly coupled network of spinal interneurons distributed throughout the spinal cord.

## Introduction

Animals produce locomotor behaviours that are adapted to environmental demands. These behaviours not only require modifications in gait but also the periodicity of movement to optimise the efficiency of locomotion. For example, long-distance migration often requires continuous locomotor movement to maximise the distance covered, whereas exploratory locomotion requires frequent pauses to survey the environment. Central pattern-generating (CPG) circuits in the spinal cord generate basic locomotor patterns, and we know that the hindlimb locomotor CPG is distributed throughout the lumbar spinal cord ^1,2^, with the rhythm generator also distributed across multiple classes of glutamatergic spinal interneuron ^3^. These glutamatergic interneurons receive sensory inputs ^4–6^, allowing for the sculpting of locomotor patterns to environmental conditions. The output of these glutamatergic interneurons is relayed to motoneurons, which in addition to controlling muscles, also activate recurrent inhibitory pathways via Renshaw cells and recurrent excitatory pathways via glutamatergic interneurons ^7,8^ in addition to other motoneurons ^9–12^ to modulate and reinforce motor output.

A key issue is that the neural circuits underlying the generation of episodic locomotor movements are poorly defined. In fish, this behaviour can be observed as a beat-and-glide movement, which emerges during early development as descending dopaminergic inputs to the spinal cord arrive ^13^ and are typically engaged during periods of exploration or foraging ^14–18^. Previous work has demonstrated that episodic locomotor patterns of activity can be generated in the developing spinal cord of fish ^14,15,18^, amphibians ^19–21^, and rodents ^22–28^. Therefore, we hypothesise that episode-generating circuits are located within the spinal cord. Whether these spinal circuits for episodic activity share properties with those generating continuous locomotion remains unclear.

To address this question, we use an *in vitro* preparation of the neonatal mouse spinal cord to examine the distribution of neural networks that underlie the genesis of episodic rhythmicity evoked by dopamine. We demonstrate that this network is distributed throughout the thoracolumbosacral spinal cord, and shares similarities yet distinct features when compared to other brainstem-spinal cord rhythms. This study provides a comprehensive analysis of episodic activity, serving as a foundation whereby future genetic and modelling ^29–32^ approaches can be used to investigate how neuromodulation can integrate with spinal circuits to produce diverse outputs.

## Results

### Episodes are flexibly coupled rostro-caudally through the spinal cord

Previous work has demonstrated the ability of multiple neuromodulators to produce patterns of episodic rhythmicity in lumbar networks studied in vitro, which are characterised by bouts of fast bursting that have an episode period of roughly 30-60 seconds ^22–26^. Here we find that episodic activity evoked by dopamine (50 µM) is not constrained to the lumbar spinal cord as it propagates to thoracic and sacral segments. Episodes of rhythmicity were recorded in ipsilateral thoracic (T9), lumbar (L1, L3, L5) and sacral (S1, S3) ventral roots (Figure 1 A; n = 7). Episodes were synchronous in 5/7 preparations (Figure 1 Bi. & C: phase = 0 -10°). In the preparations that presented with synchrony, episode onset propagated rostro-caudally (Figure 1 D; n = 5; F_(5,20)_ = 6.6, p < 0.001). This was not the case in 2/7 preparations, which presented with asynchronous episodes, particularly in the L5 and S3 segments (Figure 1 Bii. & C; phase = 90 - 180°) and is suggestive of flexible intersegmental coupling.

**Figure 1:**
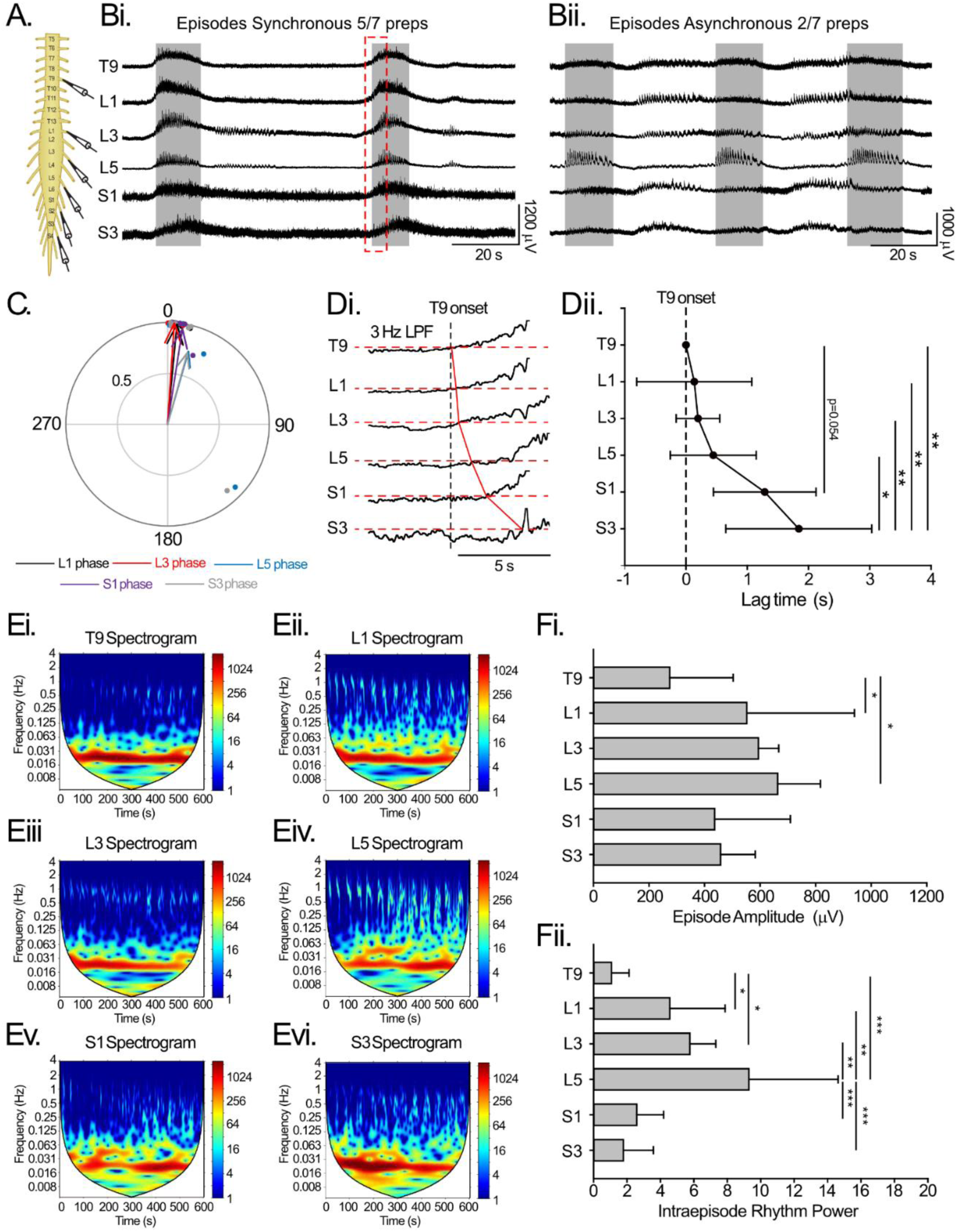
Episodes of rhythmicity are recorded in thoracic, lumbar and sacral segments. A. Suction electrodes were used to record extracellular neurograms from ventral roots of ipsilateral thoracic (T9) lumbar (L1, L3, L5) and sacral (S1, S3) spinal segments. Bi. Episodes were primarily synchronous across all spinal segments but there were instances where episodes were asynchronous (Bii.). Grey boxes are defined from episodes in the L5. The phase plot indicates that episodes were synchronous on average. C. Each point in phase plots indicates the mean vector angle and length for all episodes recorded in an individual preparation and arrows represent the group average phase angle and vector length. Di. a 3 Hz low pass filter (LPF) was applied to the region defined by a red box in Bi. to illustrate rostrocaudal propagation of episode onset for preparations that were synchronous. This is quantified in Dii with lag defined relative to episode onset in T9. E. Spectrogram analyses were performed to measure frequency power of intra-episode rhythmicity of episodes in each spinal segment where warmer colours represent higher frequency power. Fi. Episode amplitude was the largest in lumbar spinal segments and intra-episode rhythm frequency power the highest in the L5 segments (Fii). Data are presented as mean ± SD with asterisks denoting significance (*: p < 0.05, **: p < 0.01, ***: p < 0.001) from post hoc analyses following a one-way ANOVA

Episode amplitudes were highest in the lumbar segments (Figure 1 Fi; n = 7; F_(5,30)_ = 4.6, p = 0.003). The frequency power of the intra-episode rhythm was the greatest in the L5 ventral root and lower in thoracic and sacral segments (Figure 1 E & Fii; n = 7; F_(5,30)_ = 15.2, p < 0.001). There were no intersegmental differences in episode cycle periods (n = 7; F_(5,30)_ = 2.1, p = 0.1) or duration (n = 7; F_(5,30)_ = 1.9, p = 0.12) (Table 1).

**Table 1:** Parameters of episodic bursting across spinal segments. Data are presented as mean(SD) from 7 preparations.

|  | **T9** | **L1** | **L3** | **L5** | **S1** | **S3** |
| --- | --- | --- | --- | --- | --- | --- |
| Period | 47(13) | 49(14) | 55(18) | 44(15) | 48(13) | 52(7.8) |
| Duration | 24(2.5) | 32(10.2) | 36(10.6) | 29(9.2) | 27(3.4) | 28(8.8) |

### Episode-generating elements are distributed and coupled across thoracic, lumbar and sacral segments

In our next set of experiments, we set out to determine if the neural circuits that generate episodes of rhythmic activity were distributed across or localised to specific regions of the spinal cord. In contrast to episodic activity elicited by activation of sacral α_1_ adrenergic receptors or thoracic muscarinic receptors ^33^, the episodic rhythm elicited by dopamine was not dependent on phasic excitation or inhibition from segments rostral or caudal to the lumbar spinal cord. Several lines of evidence support this conclusion. First, transection of thoracic (Figure 2 Aii.; n = 7) or sacral (Figure 2 Aiii.; n = 6) spinal cord in separate experiments did not alter any parameters of episodic rhythmicity in the lumbar ventral root neurograms (Figure 2 A, D, F: duration: F_(2,17)_ = 0.8, p = 0.5; CP: F_(2,17)_ = 2.0, p = 0.2; amplitude: F_(2,17)_ = 1.8, p = 0.2; intra-episode power: F_(2,17)_ = 2.1, p = 0.15). Second, episode parameters, including cycle period, duration, amplitude and intra-episode rhythm power, were unaltered in thoracic segments following sacral transection or sacral segments following thoracic transection (Figure 2A, F). However, the coupling of episodes between T9-L5 was disrupted following transection of the sacral spinal cord (Figure 2 Biii; Phase: 36° r = 0.4 p = 0.35). This was not the case between L5 and S3 following transection of the thoracic spinal cord as the phase relationship and vector length were the same as intact preparations (Figure 2 Bii.; phase = 25° r = 0.75 p = 0.01).

**Figure 2:**
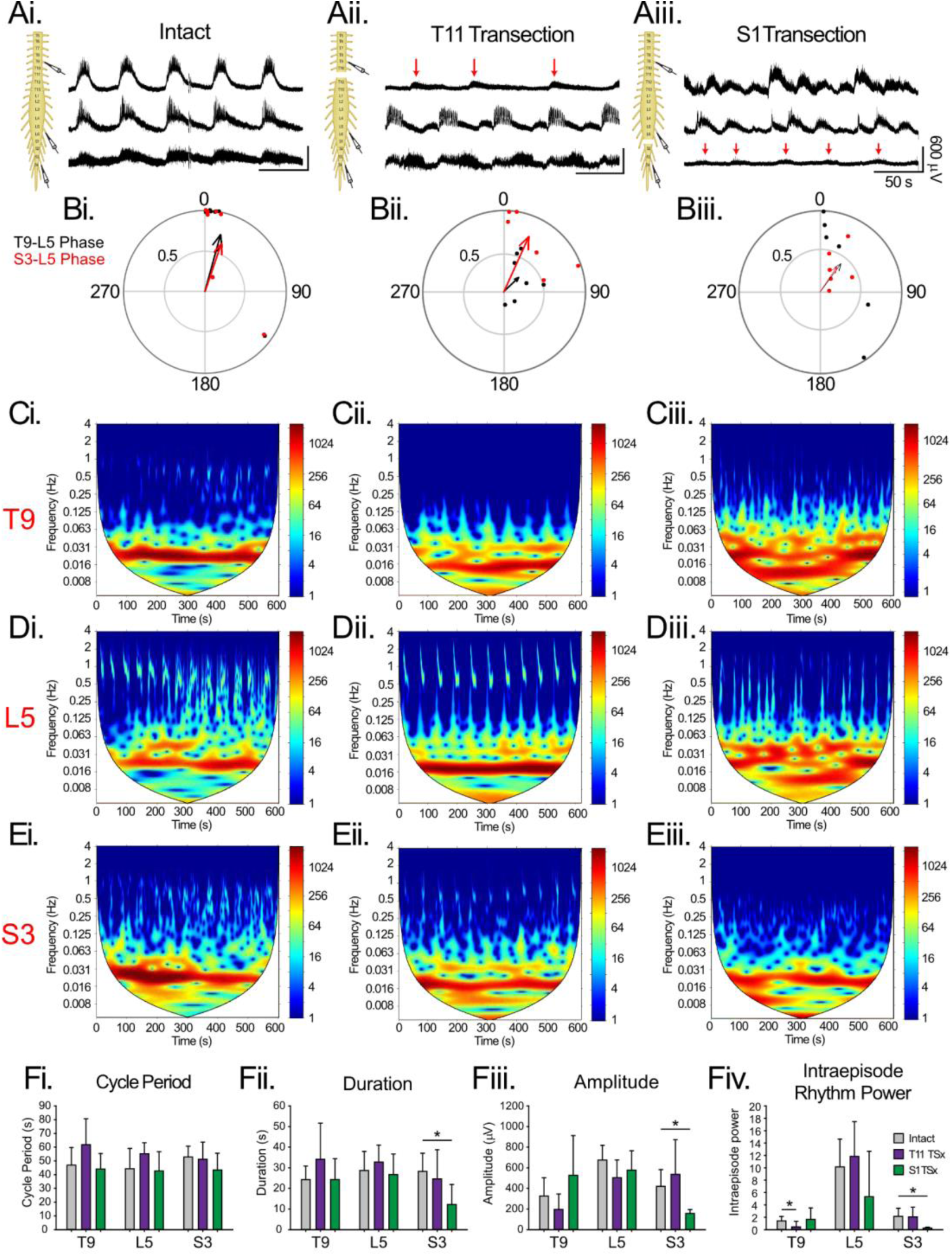
Episodes of rhythmicity evoked in isolated thoracic and sacral segments. A. Suction electrodes were used to record extracellular neurograms from ventral roots of ipsilateral thoracic (T9) lumbar (L5) and sacral (S3) spinal segments during dopamine-evoked episodic rhythms in spinal cords that were intact (Ai.), transected at the 11th thoracic segment (T11: Aii.) or first sacral segment (S1: Aiii.). Red arrows highlight low amplitude episodes detected in isolated thoracic (Aii.) or sacral (Aiii.) segments. B. The phase of episodes detected in thoracic (black) and sacral (red) segments were measured relative to those in the L5. Each point in phase plots indicates the mean vector angle and length for all episodes recorded in an individual preparation and arrows represent the group average phase angle and vector length. C-E. Spectral analyses were used to measure frequency power of intra-episode rhythmicity from episodes in thoracic (Ci-iii), lumbar (Di-iii) and sacral (Ei-iii) in preparations that were intact (Ci-Ei) transected at T11 (Cii-Eii) or S1 (Ciii-Eiii). Episode cycle period (Fi), duration (Fii), amplitude (Fiii) and intra-episode rhythm frequency power (Fiv) were measured in thoracic, lumbar and sacral ventral roots in preparations that were intact (grey bars) or transected at T11 (purple bars) or S3 (green bars). Data are presented as mean ± SD with asterisks denoting significance (*: p < 0.05, **: p < 0.01, ***: p < 0.001) post hoc analyses following a one-way ANOVA.

Episodes were also detected in isolated thoracic and sacral segments following transection. Episodes recorded in isolated thoracic segments did not differ from intact preparations in terms of cycle period (Figure 2Fi.; F_(2,17)_ = 2.8, p = 0.09), duration (Figure 2 Fii.; H_(2)_ = 3.8, p = 0.15) or amplitude (Figure 2 Fiii.; F_(2,17)_ = 2.8, p = 0.09). In the isolated sacral cord, however, episodes were shorter (Figure 2 F ii.; Duration: F_(2,17)_ = 3.56, p = 0.05) and smaller (Figure 2 Fiii.; Amplitude: H_(2)_ = 8.5, p = 0.02) but did not differ from intact preparations in the cycle period (Figure 2 Fi.; F_(2,17)_ = 1.3, p = 0.3). In both cases, episodes recorded in isolated segments did not have a superimposed fast rhythm indicated by a significant reduction in intra-episode frequency power (Figure 2 Fiv.; Thoracic: H_(2)_ = 7.9, p = 0.02; sacral: H_(2)_ = 8.9, p = 0.01). As would be expected, the coupling of episodes recorded in isolated segments and the L5 were completely disrupted, indicated by a reduction in the vector length and phase compared to episodes in the L5 (Figure 2 B). These results were reproduced following bilateral lesion of the ventrolateral funiculus at either S1 (n = 4) or T11 (n = 4), indicating that this tract is a critical pathway for coupling of episodes (Figure 3).

**Figure 3:**
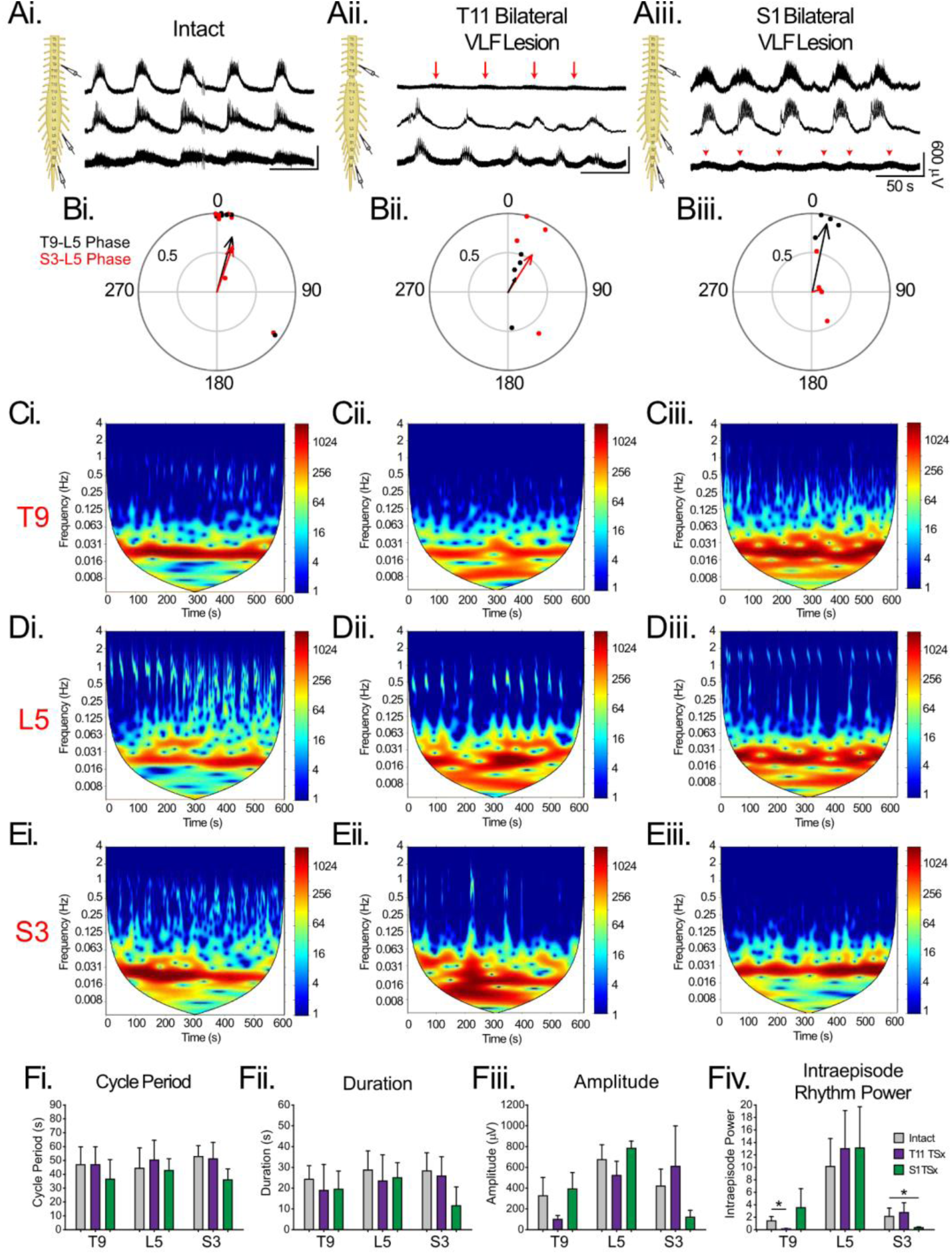
Episodes are coupled across segments through ventrolateral funiculus-projecting connections. A. Suction electrodes were used to record extracellular neurograms from ventral roots of ipsilateral thoracic (T9) lumbar (L5) and sacral (S3) spinal segments during dopamine-evoked episodic rhythms in spinal cords that were intact or following bilateral ventrolateral funiculus (VLF) lesions at the 11th thoracic segment (T11) or first sacral segment (S1). Red arrows highlight low amplitude episodes detected in thoracic (Aii.) or sacral (Aiii.) segments distal to VLF lesions. B. The phase of episodes detected in thoracic (black) and sacral (red) segments were measured relative to those in the L5. Each point in phase plots indicates the mean vector angle and length for all episodes recorded in an individual preparation and arrows represent the group average phase angle and vector length. C-E. Spectral analyses were used to measure frequency power of intra-episode rhythmicity from episodes in thoracic (Ci-iii), lumbar (Di-iii) and sacral (Ei-iii) segments in preparations that were intact (Ci.-Ei.) lesioned at T11 (Cii.-Eii.) or S1 (Ciii.-Eiii.). Episode cycle period (Fi.), duration (Fii.), amplitude (Fiii.), and intra-episode rhythm frequency power (Fiv.) were measured in thoracic, lumbar and sacral ventral roots in preparations that were intact (grey bars) or lesioned at T11 (purple bars) or S3 (green bars). Data are presented as mean ± SD with asterisks denoting significance (*: p < 0.05, **: p < 0.01, ***: p < 0.001) post hoc analyses following a one-way ANOVA.

Together, these data suggest a distributed, yet coupled oscillator with the necessary circuitry to generate the faster locomotor-like rhythm in the lumbar segments. Interestingly, the lumbar spinal cord is not necessary to generate slow episodes without superimposed rhythmicity in thoracic and sacral segments, which is in contrast to fictive locomotor activity^34^ and is more in line with the putative network interactions that generate spontaneous network activity in isolated developing spinal cords ^35–37^.

### Episodic rhythms can be entrained by sensory afferents

Information about the environment transmitted via sensory afferents is important for controlling rhythm-generating circuits and modifying movement timing and patterning in response to changing conditions. Furthermore, sensory afferents have been well documented to project across segments of the spinal cord providing a potential means of coupling episodes across segments. We therefore next determined whether the putative episode-generating oscillator was under the control of sensory afferents, as has been well-established in locomotor circuits ^38,39^, and whether they could contribute to coupling activity across segments.

Trains of electrical stimulation applied to the L5 dorsal root (Intensity: 1.5 x TH, Duration: 1 s, Train fequency: 20 Hz, Pulse width: 0.4 ms) at 1.2 x the mean dopamine-evoked episode cycle period at baseline (Figure 4 A; Rhythm cycle period at baseline: 41.2 ± 10.6 s; Stim CP: 61.3 ± 15.8 s) entrained episodes recorded in the L5 (Figure 4; n = 6). This entrainment is indicated by a reduction in episode phase variability relative to the timing of stimulation (Figure 4 B, C; Table 2), an increase in vector length (Figure 4 D) relative to the timing of stimulation, and increase in episode cycle period (Figure 4 E, F, C; n = 6; F_(2,10)_ = 11.8, p = 0.002) during the time when the stimulator was turned on. The entrainment of episodic activity in response to stimulation of the L5 dorsal root was not restricted to that particular segment and was able to entrain episodes recorded in thoracic and sacral segments (Table 2). Episodes were also entrained at an interval of 1.0 x the baseline episode period, which did not change the cycle period, but was effective in reducing the phase variance (n = 8; Table 2). Together, these results suggest that local afferent inputs can entrain the neural circuits governing episodes across spinal segments and that low-threshold afferents would be capable of triggering or modifying episodic rhythmicity.

**Figure 4:**
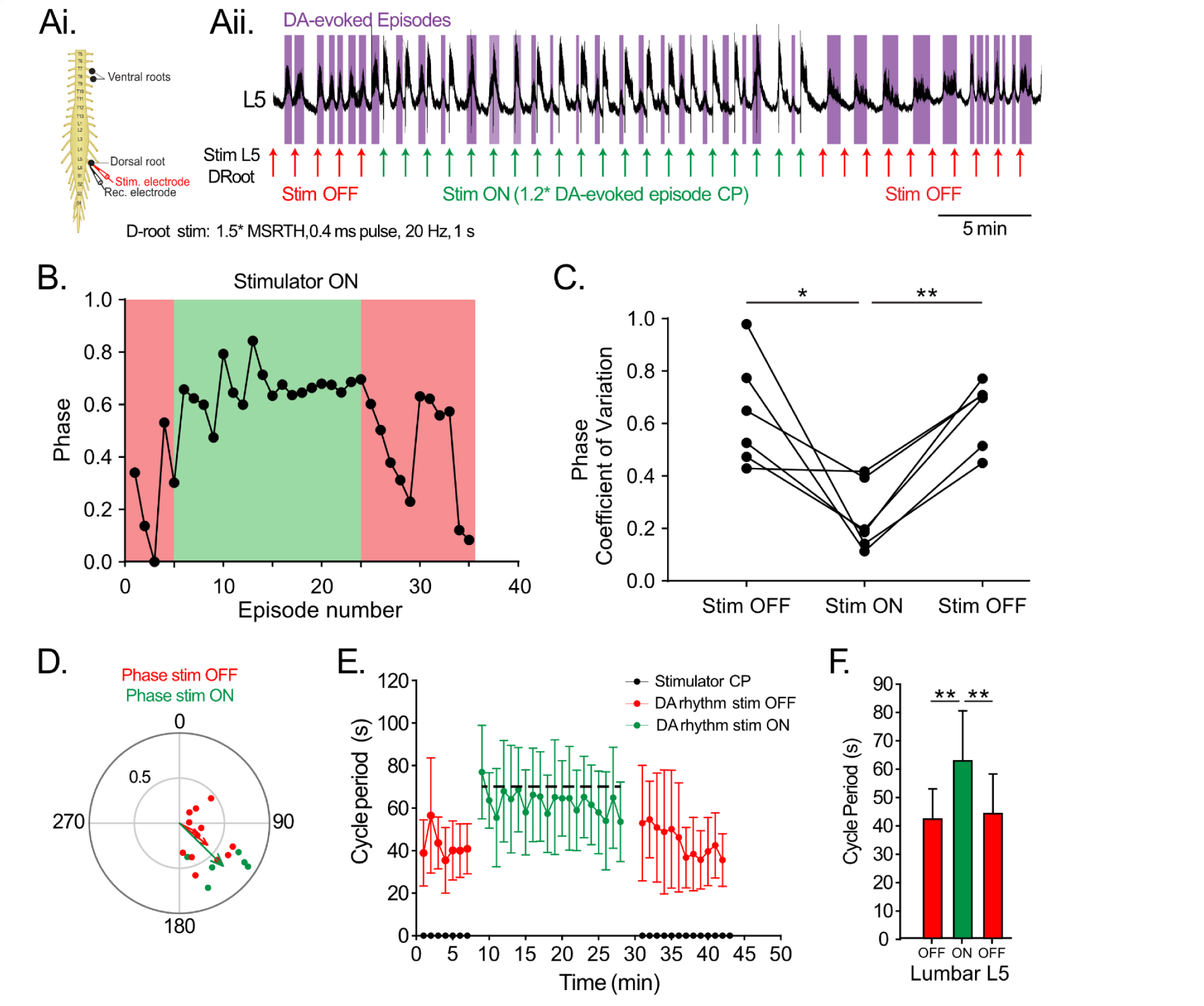
Episodes can be entrained by sensory afferents. A. Suction electrodes were used to record extracellular neurograms from ventral roots of ipsilateral thoracic (T9) lumbar (L5) and sacral (S3) spinal segments during dopamine (DA) evoked episodic rhythms. Trains of electrical stimuli were applied through a suction electrode attached to the dorsal root of the L5 (1.5xTH, 1 s, 20Hz, 0.4 ms pulse width) at 1.2 x the mean episode cycle period of the dopamine-evoked rhythm at baseline. Upward arrows represent stimulation times when the stimulator was turned off (red arrows) compared to turned ON (green arrows). Purple boxes represent spontaneously-occurring episodes evoked by dopamine in the L5. B. Representative graph plotting phase of spontaneously-occurring episodes relative to stimulation wave-marks when the stimulation was turned off (red) compared to on (green). C. Stimulation reduced the phase coefficient of variation. D. Phase plot illustrates the mean phase of dopamine-evoked episodes in lumbar relative to stimulation times when the stimulator was turned off (red) and on (green). Each point in phase plots indicates the mean vector angle and length for all episodes recorded in an individual preparation and arrows represent the group average phase angle and vector length. An increase in the vector length when the stimulator is on indicates that dopamine-evoked episodes become entrained to stimulation of the dorsal root. E. Episode cycle period increased in the L5 when the stimulator was turned on (green) and returned back to baseline when the stimulator was turned off (red). F. Episode cycle period increased when the stimulator was on (green) and returned to baseline when the stimulator was off (red bars). Data are presented as mean ± SD with asterisks denoting significance (*: p < 0.05, **: p < 0.01, ***: p < 0.001) post hoc analyses following a repeated measures one-way ANOVA within each ventral root.

**Table 2:** Episodic activity can be entrained by sensory afferents. Data are presented as mean(SD).

| **1.5 x TH, 1.2 x CP** | **N = 6** | **BL** | **Stim ON** | **Stim Off** | **F, p** |
| --- | --- | --- | --- | --- | --- |
| **Episode Cycle Period (s)** |  |  |  |  |  |
|  | T9 | 35.6(10.3) | 61.4(19.2)** | 44(15.3) | 6.5, 0.02 |
|  | L5 | 42(10.6) | 63(17)** | 44(14) | 11.8, 0.007 |
|  | S3 | 42(17.5) | 61(18.3)* | 41(9.4) | 6.5, 0.04 |
| **Episode Phase (degrees, r, p)** |  |  |  |  |  |
|  | T9 | 111, 0.34, 0.6 | 143, 0.7, 0.08 | 93, 0.38, 0.5 |  |
|  | L5 | 127, 0.37, 0.4 | 134, 0.7, 0.05 | 117, 0.26, 0.7 |  |
|  | S3 | 111, 0.34, 0.66 | 153, 0.7, 0.13 | 66, 0.38, 0.65 |  |
| **Phase Variance (CV)** |  |  |  |  |  |
|  | T9 | 0.73(0.4) | 0.23(0.06)** | 0.7(0.2) | 6.3, 0.05 |
|  | L5 | 0.64(0.2) | 0.24(0.13)** | 0.64(0.13) | 9.0, 0.02 |
|  | S3 | 0.8(0.3) | 0.25(0.06)*** | 0.8(0.07) | 13.4, 0.03 |
| **1.2 x TH, 1.0 x CP** | **N = 8** |  |  |  |  |
| **Episode Cycle Period** |  |  |  |  |  |
|  | L5 | 37(11.4) | 44(13.3) | 39(7.3) | 2.6, 0.2 |
| **Phase Variance** |  |  |  |  |  |
|  | L5 | 0.65(0.3) | 0.21(0.12)* | 0.51(0.27) | 7.9, 0.005 |

### Episodes are detected in dorsal roots but not dependent on dorsal inhibitory interneurons

Sensory afferents are integrated by interneurons located in the superficial and deep dorsal horn of the spinal cord and may therefore participate in the generation of episodic activity. Furthermore, spontaneous network activity generated by isolated spinal circuits in vitro has been reported to be partially generated through depolarization of primary afferents (PAD) by dorsal inhibitory interneurons^40^. We were interested in determining if episodes of rhythmicity evoked by dopamine were generated by similar mechanisms as spontaneous network activity. As episodic activity can be elicited by activating Drd1/5 receptors, we first performed in situ hybridization using RNAscope^®^ to examine dopamine receptor (Drd1-5) transcript expression in NeuN+ cells located in the dorsal lamina (LI-V) of the second (L2) and fifth (L5) spinal segments. A comprehensive analysis of Drd1-Drd5 receptor transcript expression across all laminae can be found in Supplementary Figure 1-3. However, to test our hypothesis, we placed particular emphasis on Drd1/5 receptor-expressing neurons that were localised to the superficial and deep dorsal laminae. In line with our hypothesis, we found subsets of NeuN+ cells that were enriched in Drd1 or Drd5 mRNA in LI-V with no observed differences between L2 and L5 segments (Figure 5 A-E; F(1,54) = 0.5, p = 0.5). Of particular note, we observed a significantly higher proportion of cells that express Drd1 (F(1,36) = 17.9, p = 0.0002) but not Drd5 receptor (F(1,18) = 2.4, p = 0.15) transcripts in dorsal laminae compared to ventral laminae (Supplementary Figure 3) supporting a potential role for dorsal circuits in the generation of episodic activity. In line with dopamine receptor expression, episodes of rhythmicity could also be detected in the dorsal roots of both the L2 and L5 segments supporting a role for PAD-generating interneurons in the generation of episodic activity (n = 6; Figure 5 F, G). Episodes recorded from the dorsal root neurograms were shorter (duration: Figure H; F_(1,5)_ = 32, p = 0.002), smaller (Table 3; F_(1,5)_ = 95.4, p = 0.0002) and degraded over the duration of an episode (Figure 5 I). Episode period did not differ between dorsal and ventral roots (Table 3; F_(1,5)_ = 0.8, p = 0.4). Overall, episodes detected in the dorsal roots were synchronous with the ventral root (Figure 5 Ji.; L2: phase = 18° r = 0.9 p = 0.01; L5: phase = 11° r = 0.96 p = 0.0008) but the onset was delayed (L2: 1.54 ± 0.46 s; L5: 1.55 ± 1.3 s) compared to episode onset in the ventral roots. Intra-episode rhythmicity in the dorsal roots was synchronous with that of the ventral root (Figure 5 Jii.; L2: phase = 51° r = 0.8 p = 0.01; L5: phase = 25° r = 0.9 p = 0.004) but restricted to the first half of an episode recorded in the lumbar ventral root neurograms. This is indicated by a significant reduction in intra-episode rhythm frequency power over time (Figure 5 I; F_(1,5)_ = 10.7, p = 0.02) and shorter episode duration in the dorsal root compared to the ventral root neurograms (Figure 5 H).

**Figure 5:**
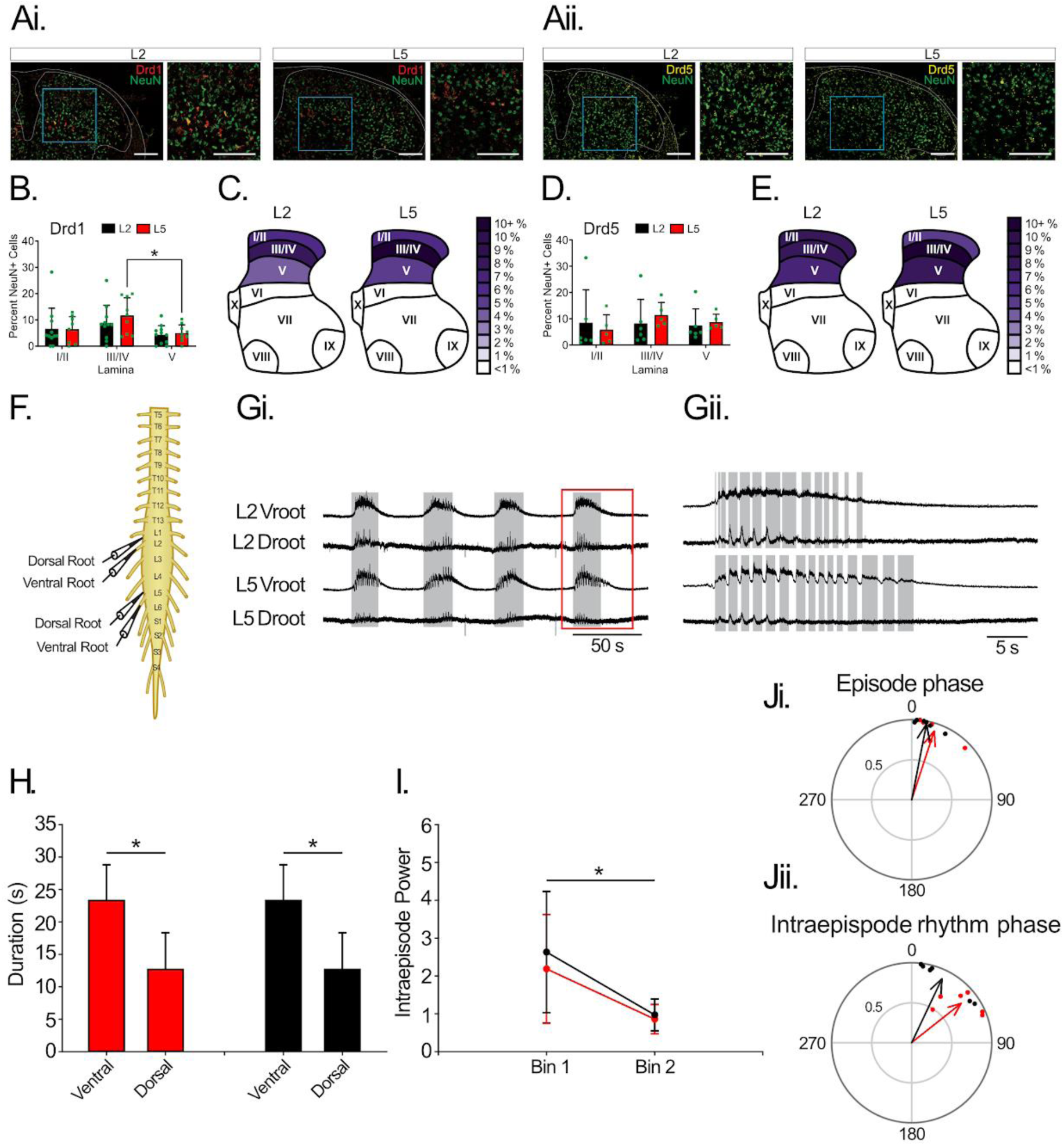
Episodes of rhythmicity recorded in dorsal and ventral roots. A. Representative image of Drd1 (Ai, red) and Drd5 (Aii, yellow) expression in NeuN+ cells (green) of dorsal lamina (LI-V) in the L2 and L5 segments of the spinal cord. Scale bars are 100 µm. B. Drd1 transcripts were expressed in a proportion of NeuN+ cells in the dorsal lamina of the L2 (black) and L5 (red) spinal segments. C. Purple heat maps depict the average expression of Drd1 transcripts in NeuN+ cells in the dorsal lamina of the L2 and L5 spinal segments. D. Drd5 transcripts were expressed in a proportion of NeuN+ cells in the dorsal lamina of the L2 (black) and L5 (red) spinal segments. E. Purple heat maps depict the average expression of Drd5 transcripts in NeuN+ cells in the dorsal lamina of the L2 and L5 spinal segments. F. Suction electrodes were used to record extracellular neurograms from ipsilateral dorsal and ventral roots of the second (L2) and fifth (L5) lumbar segments. Gi. Episodes of rhythmicity were recorded in both ventral and dorsal roots of the L2 and L5 segments. Grey boxes are defined by episodes in the L2 ventral root. Gii. Illustrates an expanded episode defined by the red box in Gi with grey boxes defined by intra-episode bursting in the L2 and L5 ventral root. H. Episodes were longer in duration in the ventral compared to dorsal roots of both the L2 (red bars) and L5 (black bars) segments. I. Frequency power of the fast rhythm measured with cross-wavelet spectral analysis declined over the duration of an episode in both the L2 (red lines) and L5 (black lines) segments. J. Phase plots for episodes (Ji.) and intra-episode (Jii.) rhythms indicate synchrony between dorsal and ventral root activities in both the L2 (red arrows) and L5 (black arrows). Each point in phase plots indicates the mean vector angle and length for all episodes recorded in an individual preparation. Data are presented as mean ± SD with asterisks denoting significance (*: p < 0.05, **: p < 0.01, ***: p < 0.001) post hoc analyses following a 2-way ANOVA.

**Table 3:** Parameters of episodic activity recorded in dorsal and ventral roots (n = 6) and effects of blocking NKCC1 pump activity that maintains presynaptic inhibition (n = 3). Data are presented as mean(SD).

| N = 6 | L2 |  | L5 |  | F, p |  |
| --- | --- | --- | --- | --- | --- | --- |
|  | Ventral Root | Dorsal Root | Ventral Root | Dorsal Root |  |  |
| Period | 41(5.6) | 38(3.6) | 39(12) | 39(15) | 0.8, 0.4 |  |
| Amplitude | 817(450) | 144(55) | 841(500) | 77(20) | 95, 0.002 |  |
| N = 3 | Dorsal Roots |  |  | Ventral Roots |  |  |
|  | DA | DA+Bumetanide | T(6) | DA | DA+Bumetanide | T(7), p |
| Period | 42(10.8) | 39(2.6) | 0.4, 0.7 | 43(11.6) | 46(8.2) | 0.4, 0.7 |
| Duration | 13.3(6.5) | 12.1(14) | 0.2, 0.9 | 23(6.5) | 26(4.5) | 0.8, 0.5 |
| Amplitude | 106(43) | 132(35) | 0.8, 0.4 | 837(281) | 976(255) | 0.7, 0.5 |

As episodic activity could be detected in dorsal roots, we next tested whether dorsal circuits that produce primary afferent depolarization are involved in the generation of episodic activity. To test this hypothesis, we bath applied the NKCC1 inhibitor (50 µM; n = 3), bumetanide, to disrupt the depolarizing influence of presynaptic inhibitory interneurons onto primary afferents in the dorsal horn^40^. However, episodes recorded in the dorsal and ventral roots were unaltered by bumetanide (Table 3), suggesting that episodes are not generated by the depolarization of primary afferents through dorsal inhibitory neuronal populations.

### Episodic activity is generated by ventral interneurons and not recurrent pathways

Locomotor-like activity is generated by neurons located in the ventral horn of the spinal cord with emerging roles for recurrent pathways formed between motoneurons and premotor inputs being revealed more recently ^9,10,12,41,42^. We therefore hypothesised that Drd1/5 receptors may produce slow oscillations in motoneurons or premotor interneurons leading to the generation of episodic activity as has been reported in deep dorsal interneurons ^43^. In line with our hypothesis, we identified a small proportion of neurons in Laminas VII, VIII, and IX that are enriched in Drd1 and Drd5 mRNA, with no differences observed between L2 and L5 segments (Figure 6 A-E; F(1,54) = 1.7, p = 0.2).

**Figure 6:**
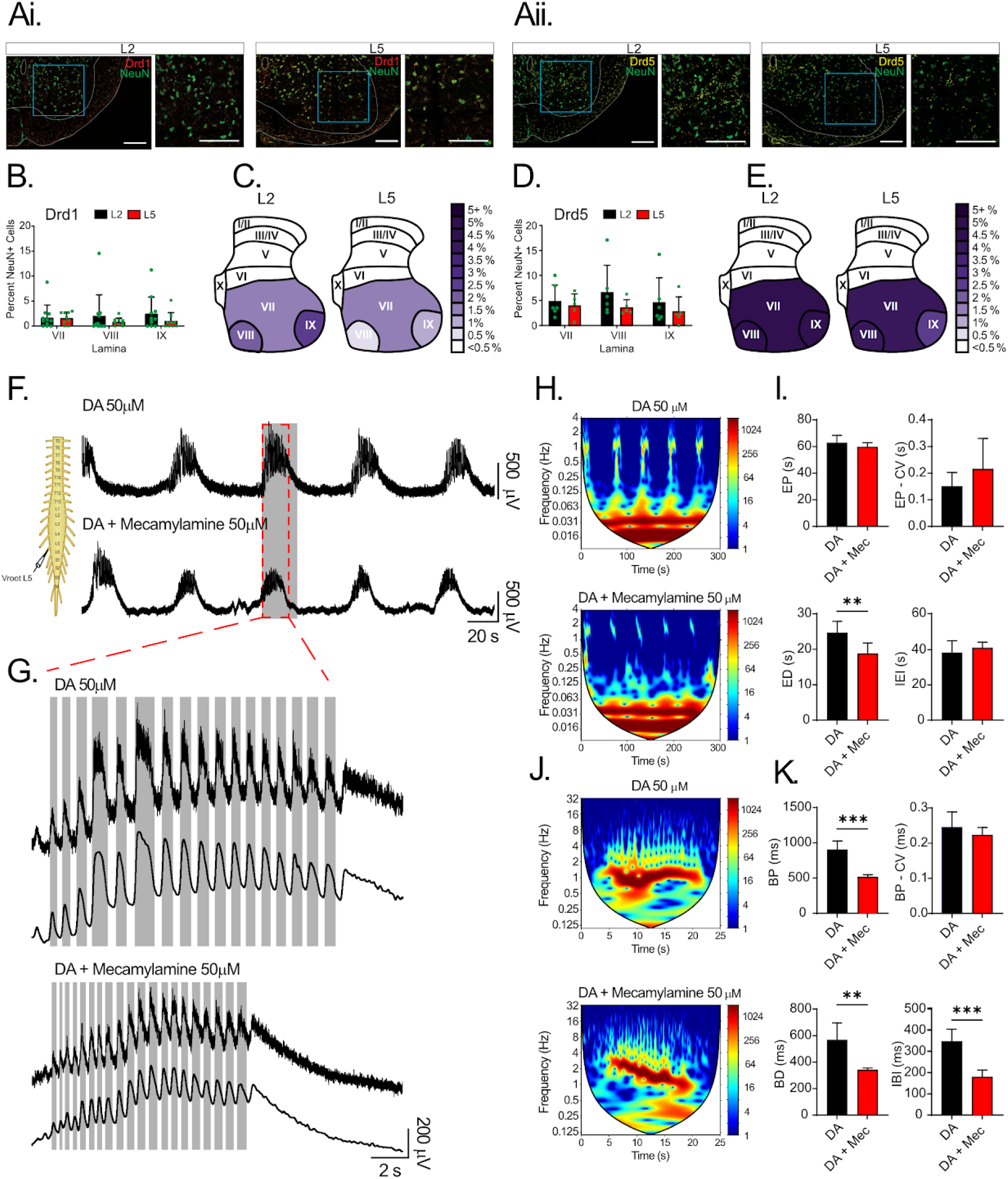
Episodes of rhythmicity are affected but not mediated by cholinergic pathways. A. Representative image of Drd1 (Ai, red) and Drd5 (Aii, yellow) expression in NeuN+ cells (green) of ventral lamina (LVII - IX) in the L2 and L5 segments of the spinal cord. Scale bars are 100 µm. B. Drd1 transcripts were expressed in a proportion of NeuN+ cells in the ventral lamina of the L2 (black) and L5 (red) spinal segments. C. Purple heat maps depict the average expression of Drd1 transcripts in NeuN+ cells in the ventral lamina of the L2 and L5 spinal segments. D. Drd5 transcripts were expressed in a proportion of NeuN+ cells in the ventral lamina of the L2 (black) and L5 (red) spinal segments. E. Purple heat maps depict the average expression of Drd5 transcripts in NeuN+ cells in the ventral lamina of the L2 and L5 spinal segments. F. Suction electrodes were used to record extracellular neurograms from the ventral root of ipsilateral lumbar (L5) spinal segment during dopamine-evoked episodic rhythms; recordings show the effects of dopamine alone and after application of Mecamylamine. G. Display of expanded episodes defined as a red dashed box illustrating raw (top) and low pass filtered (3 Hz, bottom) traces. Grey boxes define intra-episode bursting in the ventral root. H. Spectrogram analyses from episodes recorded in ventral ventral root L5. I. Analysis of the episode cycle period, coefficient of variation, duration, and inter-episode interval, measured in the lumbar ventral root L5 during dopamine-evoked (red) episodic rhythms and after Mecamylamine (black). J. Spectrograms from the intra-episode rhythm in G. K. Same as in I. for the intra-episode rhythm analysis. Data are presented as mean ± SD with asterisks denoting significance (**: p < 0.01, ***: p < 0.001) from post hoc analyses following an unpaired *t*-test.

We next set out to determine whether slow oscillations in motoneurons could give rise to episodic activity through recurrent pathways^10^. To test this hypothesis, we applied the nicotinic receptor antagonist mecamylamine (50 uM) to block cholinergic signalling acting through ionotropic nicotinic receptors, which are important for the activation of Renshaw cells that produce recurrent inhibition. In contrast to our hypothesis, mecamylamine produced only modest effects on dopamine-evoked episodes, reducing episode duration (t_(5)_ = 4.1, p = 0.009) with no effect on episode period (t_(5)_ = 1, p = 0.325), variation (t_(5)_ = 1.6, p = 0.163), interepisode interval (t_(5)_ = 0.9, p = 0.381), or amplitude (t_(5)_ = 1.6, p = 0.159; Figure 6 F-I). Interestingly, we found that mecamylamine produced robust effects on the faster intra-episode rhythm. When applied during stable episodic activity, mecamylamine decreased the intra-episode burst period (t_(8)_ = 7.6, p < 0.001), duration (t_(8)_ = 4.4, p = 0.002), and inter-burst interval (t_(8)_ = 5.9, p < 0.001), but did not alter variance (t_(8)_ = 1, p < 0.309) or amplitude (t_(8)_ = 0.6, p = 0.549; Figure 6 G, J, K). These results suggest a role for nicotinic acetylcholine receptors in the control of the fast intra-episode (locomotor) rhythm but does not support a role for recurrent pathways acting through Renshaw cells in the generation of episodic activity.

Motoneurons also provide glutamatergic recurrent excitation of other motoneurons ^9,11^ and V3 interneurons ^7^, which may also contribute to the generation of episodic activity. To test this possibility, we recorded the activity of premotor interneurons that project through the ventrolateral funiculus (VLF) adjacent to motoneurons that project through the neighbouring ventral root (n = 4; Figure 7). If recurrent pathways contribute to the generation of episodic activity then we would expect to see episodes in both the ventral root and VLF with those in the ventral root preceding that of the VLF. In line with this possibility, episodes were recorded in the VLF, which were of similar duration (t_(3)_ = 1.8, p = 0.2) and cycle period (t_(3)_ = 1.9, p = 0.15) as the ventral root (Table 4). Episode amplitude was attenuated in the VLF compared to ventral roots (t_(3)_ = 7.8, p = 0.004) ^44^, likely reflecting a fewer axons and a lower firing rate of interneurons compared to motoneurons. Episodes recorded in the VLF were synchronous with those recorded in the ventral root (Figure 7 Ei.: Phase: 18 °, r = 0.9, Rayleigh test p = 0.02), however, in contrast to what would be expected if recurrent pathways contributed to episode generation, we found that episodes in the VLF preceded those in the ventral root by 0.97 ± 0.28 s (Figure 7 B). Episodes recorded in the VLF also displayed a superimposed fast rhythm that slowed down over the duration of an episode (VLF fast rhythm bin 1: 1.1 ± 0.2 Hz; Bin 2: 0.87 ± 0.22 Hz; t_(3)_ = 4.0, p = 0.03) similar to what we have observed in the ventral roots. Cross-wavelet analysis of intra-episode rhythmicity between the L3 VLF and L2 ventral root revealed that the intra-episode bursting in the VLF was synchronous with that of the adjacent ventral root and maintained this pattern throughout the duration of episodes (Figure 7 Eii. Vroot-VLF phase bin 1: 50 °, r = 0.63, Rayleigh test p = 0.2; phase bin 2: 57 °, r = 0.42, Rayleigh test p = 0.5). Together, these data suggest that episodes of dopamine-evoked rhythmicity are produced directly by premotor interneurons while there is a lack of support for the involvement of motoneuron-mediated recurrent excitation.

**Figure 7:**
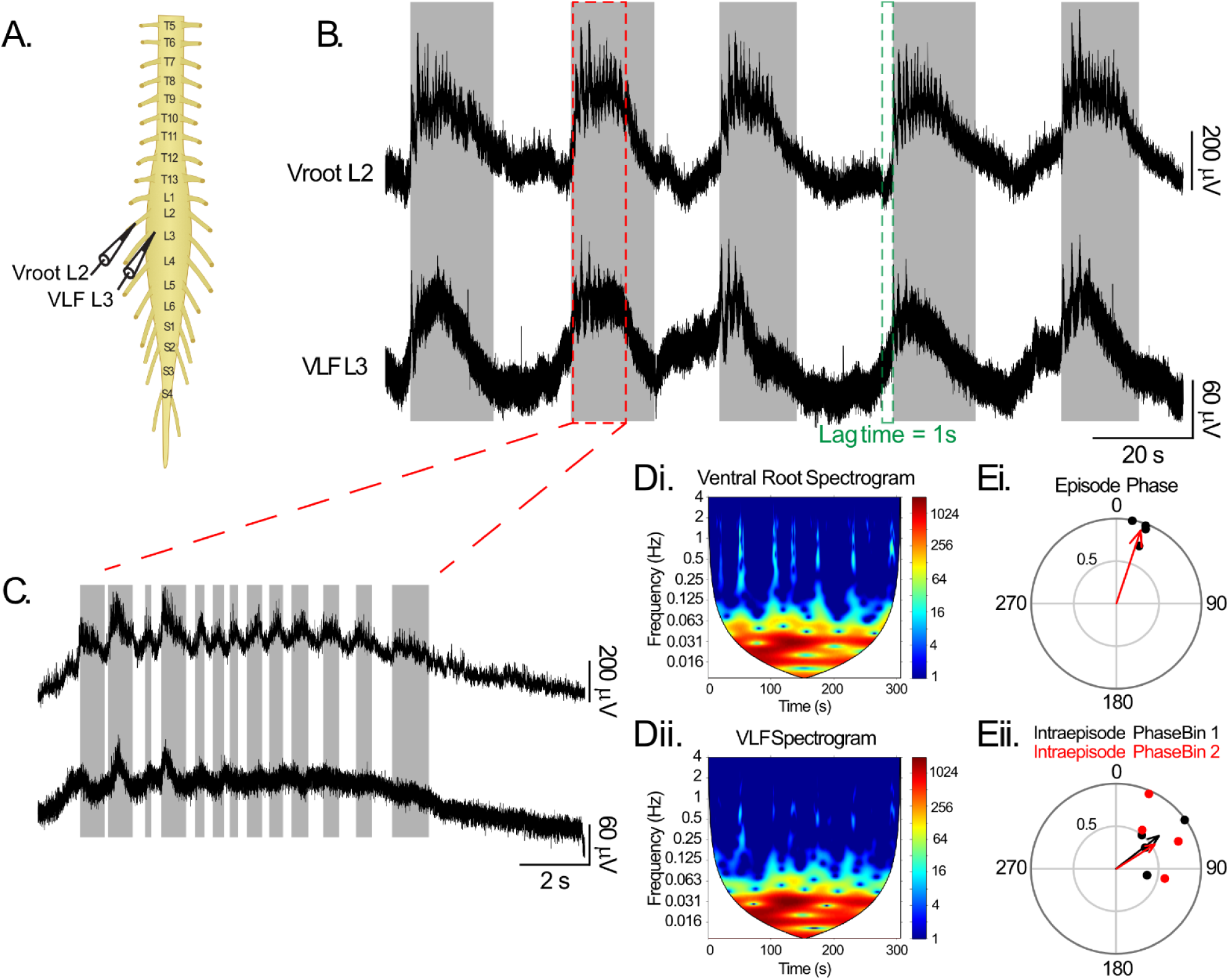
Episodes of rhythmicity recorded in ventral roots and the ventrolateral funiculus. A. Suction electrodes were used to record extracellular neurograms from neighbouring ventral roots in the L2 segment and a slip of the ventrolateral funiculus (VLF) in the L3. B. Episodes of rhythmicity were recorded in both the ventral root and VLF. Grey boxes define onsets for the ventral root episodes. The green line indicates that episode onset in the VLF precedes the ventral root. C. Display of an expanded episode defined as a red dashed box in B. to illustrate patterning of the intra-episode rhythm. Grey boxes define intra-episode bursting in the ventral root. The intra-episode rhythm was analysed using spectral analysis. D. Depicts spectrograms from episodes recorded in the ventral root and VLF. E. Phase plots illustrate episodes in the ventral root and VLF are synchronous with intra-episode rhythm also predominantly synchronous over the duration of episodes. Each point in phase plots indicates the mean vector angle and length for all episodes recorded in an individual preparation.

**Table 4:**
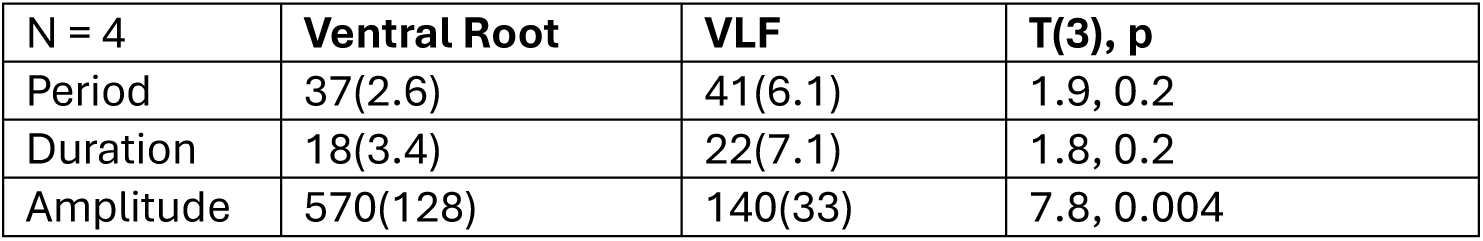
Parameters of episodic activity recorded in the ventrolateral funiculus (n = 4). Data are presented as mean(SD).

## Discussion

The rhythmic output generated by mammalian spinal networks’ to produce locomotion are commonly studied through continuous locomotor-like patterns, but natural locomotor behaviours are episodic, characterised by movement interspersed with pauses. An unresolved question is whether these episode generating circuits in the spinal cord share similar properties to those that generate continuous patterns of locomotor activity that produce the fundamental rhythm and pattern of walking. While swimming episodes in isolated spinal circuits of fish and tadpoles have been explored ^18,45,46^, the emphasis has been on understanding intra-episode features, rather than the inter episode rhythmicity ^15–17,21^. Neuromodulators can induce episodic rhythms in isolated mouse spinal cords ^22–26^ and manipulating network excitability ^24^ or modulating h-currents (regulating episode duration) and sodium-potassium pump currents (controlling inter episode intervals) can transition episodic bursting to continuous bursting ^26^. Here we deployed episodic activity elicited by dopamine in isolated neonatal mouse spinal cord preparations to determine the organisation of spinal circuits that generate episodic activity (Figure 8). Our data suggests that episodic rhythms evoked by dopamine share similarities yet distinct features from other rhythms generated by spinal networks of the neonatal rodent in vitro.

**Figure 8:**
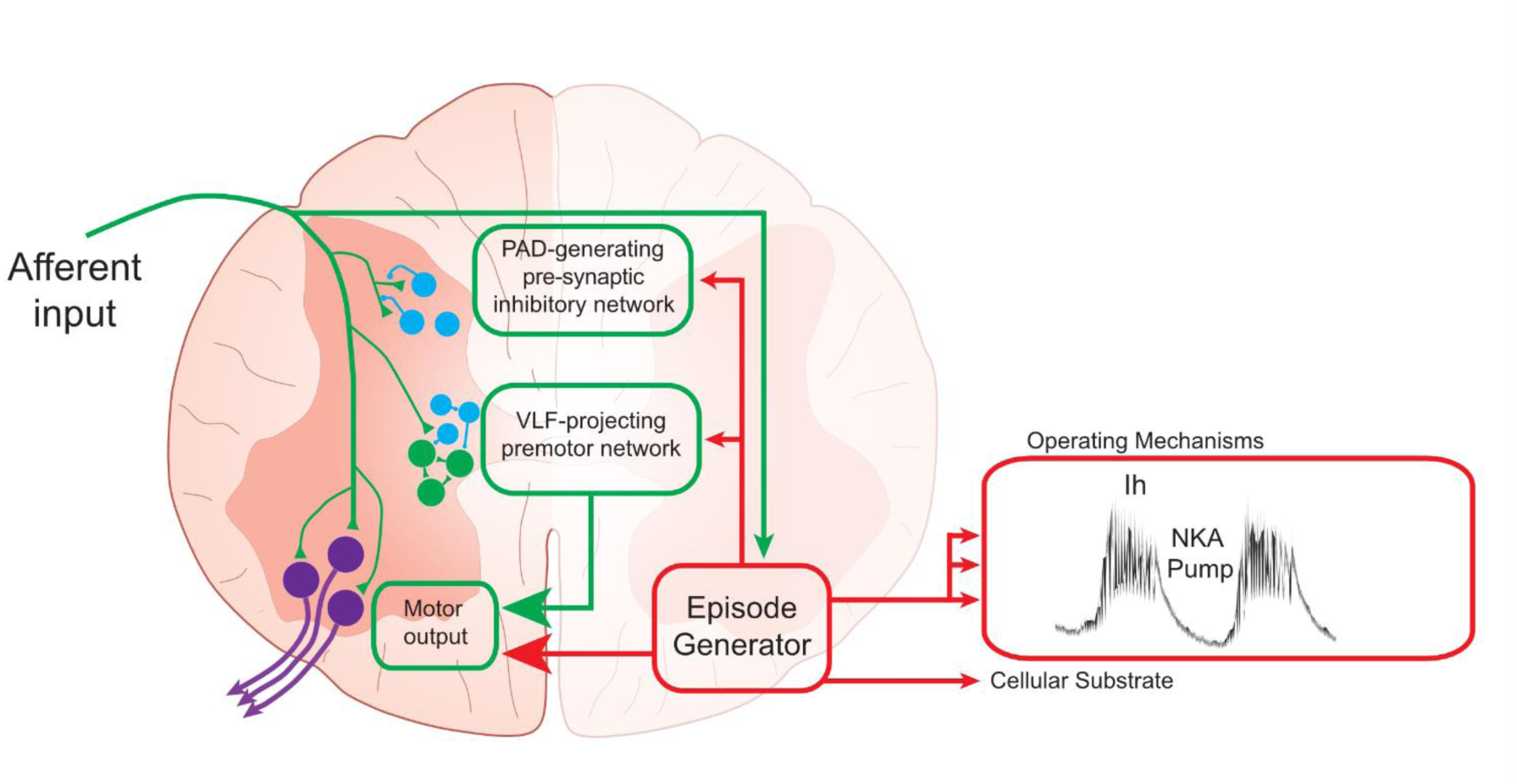
Summary Schematic.

First, there are key differences regarding spatial and temporal patterning. Segmental differences are less apparent in the patterning of the episodic rhythm compared to the locomotor rhythm. Although both locomotor-like and episodic rhythms can be recorded in thoracic, lumbar, and sacral segments^35^, they differ with respect to the direction of propagation through the spinal cord. First, dopamine-evoked episodes propagate rostro-caudally much like episodes of spontaneous activity^36^ or disinhibited bursting^34^ whereas locomotor bursts propagate caudo-rostrally^35^. Second, episodes of activity seem to be generated by a network that is distributed across thoracic, lumbar and sacral spinal segments, given that slow episodes could be generated in isolated sacral or thoracic segments following full transection or bilateral lesion of the ventrolateral funiculus. Interestingly, fast intra-episode bursting was abolished in isolated sacral or thoracic segments following transection or lesion suggesting that the distributed episode-generating network must be upstream and activate lumbar circuits that generate the fast rhythm (Figure 8). This result is in line with previous reports demonstrating the capacity of isolated lumbar segments to produce locomotor bursting ^1,2^, however, the lumbar cord is necessary for the generation of fast locomotor-like bursting in both thoracic and sacral segments. Interestingly, and in contrast to episodic activity produced by dopamine, sacral segments also display the capacity to produce continuous alternating bursting when isolated from the lumbar spinal cord ^47,48^, indicating that episodic activity elicited by dopamine might act through distinct circuits compared to those activated by other modulators in the sacral spinal cord.

Sensory circuits can also generate network activities produced in the isolated spinal cord ^49^ and through intersegmental projections, may act to couple activity across segments. Furthermore, a key feature of rhythm generating networks is the ability to entrain the rhythm or pattern by afferent inputs ^50–53^. In the spinal cord, sensory afferents appear to reset the episodic and fictive-locomotor rhythm in different ways. In the case of episodic activity, resetting through stimulation of dorsal roots in the L5 was synchronous across all spinal segments. However, for fictive locomotion, perinatally, the rhythm shows an asynchronous reset between ipsilateral L2 and L5 roots ^54^. This observation is consistent with ipsilateral inhibitory connections for the fictive locomotor circuit and potentially ipsilateral excitatory connections for the episodic network.

Having identified that sensory afferents can regulate episodic activity, we next set out to determine whether interneurons located in the dorsal horn participate in the generation of episodes. In line with our hypothesis, we found a subset of neurons in the dorsal laminae that were enriched in transcripts for Drd1 and Drd5 receptors. Furthermore, both the episode and superimposed fast rhythms can be recorded in dorsal roots, albeit the fast rhythm is restricted to the first half of the episode when fast bursting tends to be synchronous^55^ suggesting a potential role for dorsal inhibitory interneurons that contribute to depolarization of primary afferents in the generation of episodic activity. During alternating rhythmic activity, dorsal circuits appear to be de-recruited ^49^, which may be suggestive of a phase-dependent reduction in primary afferent depolarization over the duration of an episode. However, further experiments revealed that episodes are not generated by inhibitory presynaptic interneurons mediating primary afferent depolarization. Our findings demonstrate that episode onset in the ventral root precedes that in the dorsal root, and episodes can still be evoked in the presence of the NKCC1 blocker, bumetanide, which would be expected to reverse the depolarizing influence of chloride on primary afferent terminals^40^. Together these results suggest that dorsal circuits are recruited during episodic activity, however, they are not responsible for the generation of the underlying episode.

In our next set of experiments, we set out to determine whether ventral circuits produce episodic activity. Growing evidence supports roles for recurrent pathways produced by motoneurons in the modulation of locomotor network activity ^7,42^. Classically, this role was demonstrated in isolated mouse spinal cords through electrical stimulation of the ventral root as a means of eliciting locomotor-like activity ^56,57^ and has been demonstrated more recently using optogenetic activation of motoneurons ^42^. The best described of these pathways is the recurrent inhibitory pathway formed by motoneurons providing nicotinic cholinergic excitation of Renshaw cells ^58^, which in turn produces glycinergic inhibition of motoneurons ^59^. We asked whether slow oscillations in motoneurons could produce episodic activity through the engagement of cholinergic recurrent pathways ^10^. Consistent with reducing the recurrent inhibitory pathway, nicotinic acetylcholine receptor antagonists produced an increase in the frequency of the fast intra-episode rhythm. This finding is in contrast to those reported using continuous locomotor-like activity elicited by 5HT, NMDA and dopamine, where mecamylamine did not alter rhythm parameters ^60^. One possible explanation is that roles for nicotinic pathways may be diminished during the high-conductance state produced by the application of these locomotor drugs, which may have been revealed during a lower-excitability state produced during dopamine-evoked episodic bursting ^24^. Interestingly, and in contrast to our hypothesis, we found that antagonism of nicotinic receptors had minor effects on the episodic inter-episode rhythm, with only episode duration being reduced. This result suggests that undefined cholinergic interneurons may produce excitatory actions that prolong episode duration. V0c interneurons form a prominent source of cholinergic input to motoneurons, however are unlikely to mediate control of episodic activity given that they exert their influence primarily through muscarinic receptors ^61–63^.

Motoneurons also give rise to excitatory glutamatergic recurrent pathways that connect with other motoneurons ^9,11,12^ and glutamatergic V3 interneurons ^7^, which could also contribute to the generation of underlying episodes ^60,64^. However, in contrast to this hypothesis, we found the underlying slow episodes recorded in tracts projecting through the ventrolateral funiculus preceded that of motoneurons in the ventral root suggesting that episodes are likely produced by premotor interneurons. Furthermore, previous work has also demonstrated that dopamine produces Drd2-receptor-mediated inhibition of locomotor-like activity elicited by electrical stimulation of the ventral roots ^56^, which may occur through the inhibition of V3 interneurons^25^. Together, these results support a role for nicotinic pathways in the control of locomotor-like intra-episode activity; however, our data do not support the idea that recurrent pathways produced by motoneurons are responsible for the genesis of the underlying episode. Our results suggest that episodic activity is generated by premotor interneurons that are upstream from those that generate the locomotor-like rhythm (Figure 8).

We performed RNAscope^®^ in our attempt to determine the spatial localization of neurons that might generate episodic activity. Given that Drd1/5 agonists are capable of eliciting episodic activity ^25^, we hypothesised that neurons that are important for their generation would demonstrate a spatial organisation that could provide insight into functionally defined subsets as has been extensively demonstrated with the direct and indirect pathways in the striatum ^65^. Consistent with the Allen Spinal cord Atlas and previous reports ^66,67^, we found that Drd2 receptor transcripts were expressed in the most abundance and were particularly enriched in dorsal laminae. A similar observation was also made for Drd1/5 receptors leading us to test roles for dorsal circuits in the generation of episodic activity. However, as discussed previously, our results did not support this hypothesis. Interestingly, and in contrast to our hypothesis, overall, Drd1/5 receptor transcript labelling was relatively sparse in the ventral horn, suggesting that activation of a small proportion of spinal neurons by neuromodulatory pathways could have profound effects on network output. It is possible that the equally subtle differences in receptor expression in different laminae could have profound effects on output. Our results do not rule out the possibility that functionally defined subsets of spinal neurons could be identified based on neuromodulatory receptors given that functionally defined neurons in the striatum are also spatially intermingled. These results are in line with our previous reports using single cell recordings and Drd2 agonists, which suggested that Drd2-expressing interneurons are also spatially intermingled and might be expressed across multiple subsets of genetically defined populations of spinal interneurons. In all likelihood, no one class of interneuron would be responsible for the generation of a particular rhythm. Indeed there are examples from the leech^68^, tadpole^69^, and turtle^70^ where interneurons are active and contribute to multiple rhythmic movements. In the mammalian spinal cord, V3 interneurons ^71–74^ and ventral spinocerebellar tract-projecting neurons ^75^ are feasible targets given that they can both elicit episode of locomotor activity and demonstrate connectivity and intrinsic properties that are important for the generation of episodic activity. Multifunctionality within these defined populations of rhythm generating circuits therefore serve as an important direction for future study.

### Developmental considerations

We have leveraged the tractability of the neonatal mouse spinal cord to dissect the neural networks that generate rhythmic activities that underlie complex movements. It is important to recognize that these experiments were performed using a developing model system with ongoing maturation of circuits and behaviour ^76–78^. It could be argued that the episodic activities that we report in vitro may not pertain to the function of the fully mature animal. However, locomotor behaviours in adult animals are episodic in nature and episodic locomotor activities can also be readily evoked in isolated spinal circuits obtained from mature animals ^27,79,80^. This work highlights the need to better understand neural networks that encode this important feature of locomotor behaviours, alongside aspects of rhythm, pattern, and gain.

## Conclusion

Here we provide a foundational examination defining the distribution of a network that contributes to the generation of episodic rhythmicity in the developing mouse spinal cord. We expect the study of these stable non-locomotor rhythms will provide insight into mechanisms that contribute to the generation of diverse rhythmic activities in the mammalian spinal cord. This would include not only the conductances and cellular mechanisms ^14,26,27^ but also insight into how the network is reconfigured to switch between states and generate different rhythmic outputs.

## Acknowledgements

We would like to acknowledge support from the Whelan Lab, Spinal Core software was a kind gift from Prof A. Lev-Tov (Hebrew University).

## Declaration of generative AI and AI-assisted technologies

During the preparation of this work, the authors used ChatGPT to proofread for grammatical errors and reduce the abstract to 150 words. After using this tool, the authors reviewed and edited the content as needed and take full responsibility for the content of this publication.

## Contributions

Conceptualization, SAS, JJMC, PJW; Methodology, SAS, JJMC, MAT, PJW; Investigation, SAS, JJMC, APL, MAT, SADV, AS; Formal Analysis, SAS, JJMC, MAT, CL, AS, MJB; Writing - Original Draft, SAS, JJMC, PJW; Writing - Reviewing and Editing, All Authors; Funding Acquisition, PJW, SAS; Supervision, SAS, JJMC, GBM, PJW.

---

**Supplementary Figure 1:**
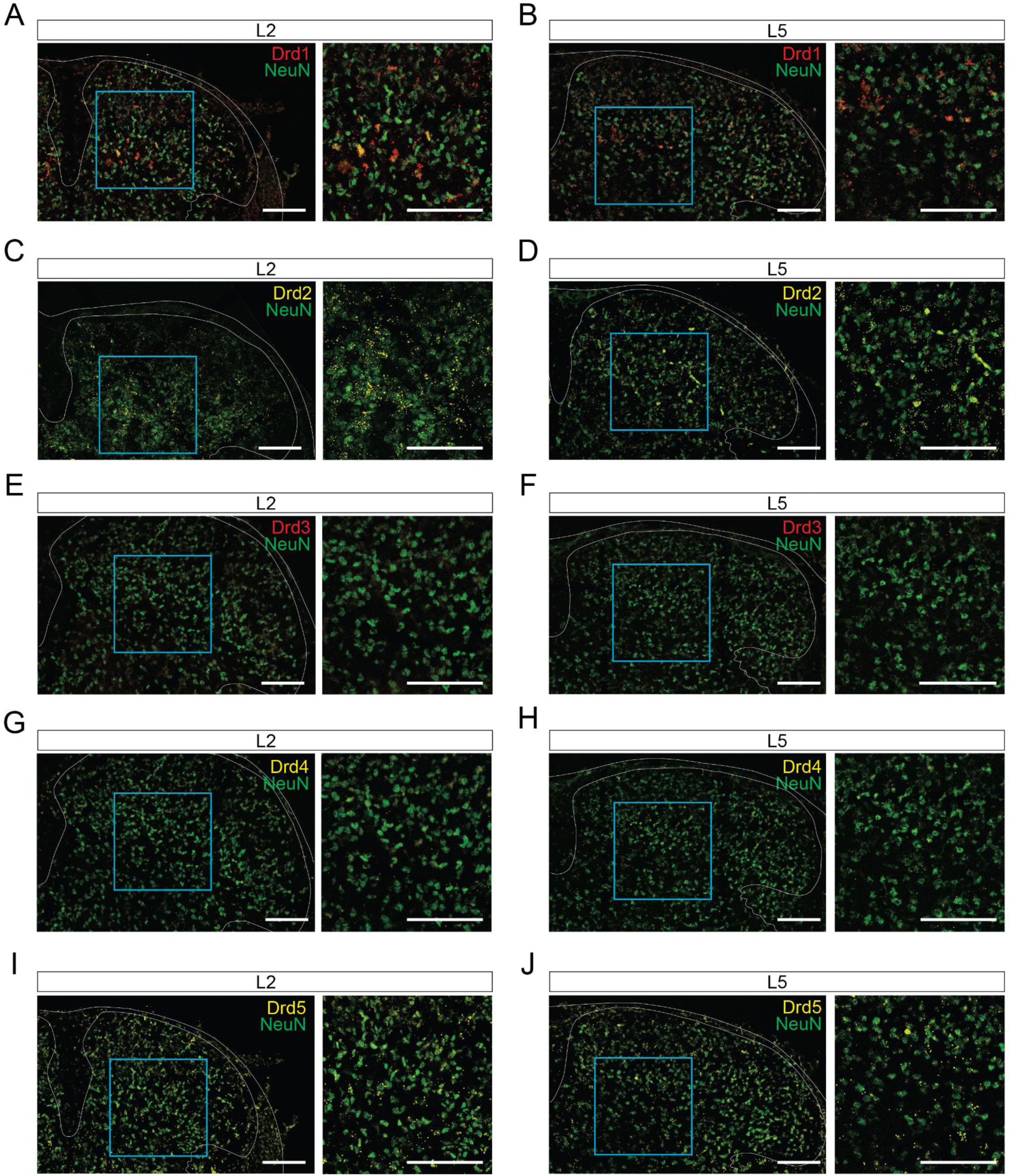
Dopamine receptor transcripts distribution through the dorsal lamina. A-J. Representative images of Drd1-5 expression in NeuN+ cells (green) of dorsal lamina (I - V) in the L2 and L5 segments of the spinal cord. Scale bars are 100 µm.

**Supplementary Figure 2:**
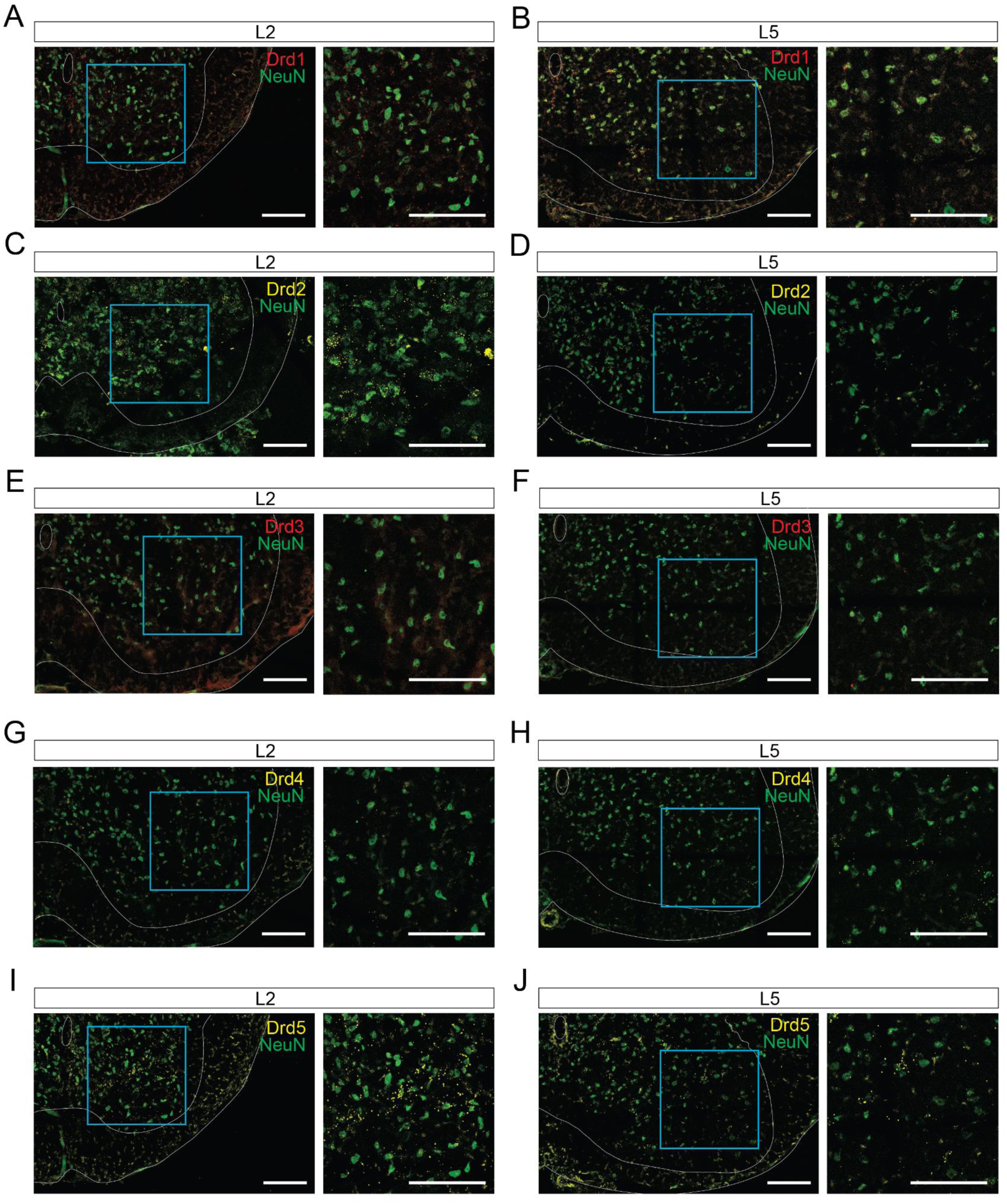
Dopamine receptor transcripts distribution through the ventral lamina. A-J. Representative images of Drd1-5 expression in NeuN+ cells (green) of ventral lamina (VII - IX) in the L2 and L5 segments of the spinal cord. Scale bars are 100 µm.

**Supplementary Figure 3:**
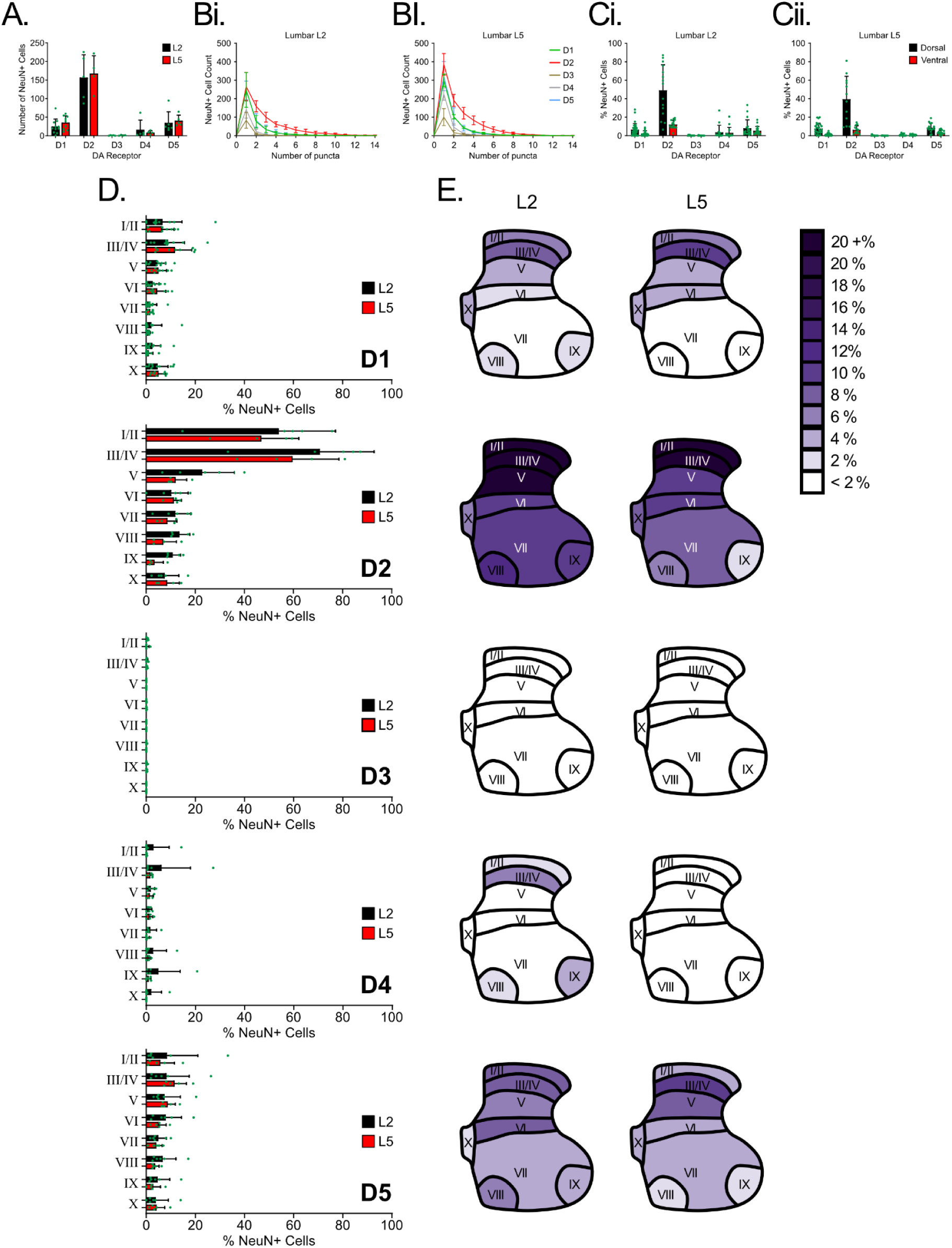
Dopamine receptor transcripts are expressed in subsets of spinal neurons. A. Total number of neurons identified by the expression of NeuN expressing dopamine receptor transcripts in the L2 (black) and L5 (red) spinal segments. B. Number of puncta detected for each dopamine receptor transcript (Drd1-5) within a region of interest defined by NeuN labelling in the L2 (Bi) and L5 (Bii) segments of the spinal cord. C. Dopamine receptor transcripts were expressed in the largest proportion of cells in the dorsal horn. D. Proportion of cells that express transcripts for each dopamine receptor across spinal laminae in the L2 (black) and L5 (red) spinal segments. E. Purple heat maps depict the average expression of Drd1-5 transcripts in NeuN+ cells through all the lamina in the L2 and L5 spinal segments. Data are presented as mean ± SD with individual data points for each animal displayed. Percentage values are expressed relative to the number of NeuN+ cells within each lamina.

## Methods

### Ethical approval & animals

Experiments were performed on neonatal C57BL/6J (N = 67) mice of postnatal 0-4 days old (P0-P4) and either sex. All procedures used were approved by the University of Calgary Health Sciences Animal Care Committee (Protocol Number: AC16-0182).

### Tissue preparation

Animals were anaesthetised via hypothermia by placing them at -20°C for 5-10 minutes. Animals were then decapitated, eviscerated and spinal column pinned ventral side up in a silicone elastomer (Sylgard) - lined dissecting dish that was superfused with room temperature (21°C) carbogenatedartificial cerebrospinal fluid (aCSF) (in mM: 4 KCl, 128 NaCl, 1 MgSO_4_, 1.5 CaCl_2_, 0.5 Na_2_HPO_4_, 21 NaHCO_3_, 30 ᴅ-glucose, 310-315 mOsm.) bubbled with carbogen (95% O_2_-5% CO_2_). The spinal cord was isolated by performing a ventral laminectomy. The nerve roots connecting the spinal cord to the vertebral column were subsequently cut, isolating the spinal cord. The isolated spinal cord was then transferred to a recording chamber perfused with aCSF bubbled with carbogen and placed ventral side up. The bath temperature was then gradually increased to 27°C (Whelan et al., 2000), the advantage of this is that it is closer to physiological temperatures and avoids fluctuations in room temperature. Spinal cords were left to stabilise for 1 hour before experiments were performed. For RNAscope^®^ experiments, the spinal cord was isolated and immediately frozen in dry ice. Tissue was placed in cryomolds and frozen in Optimal cutting tissue medium (OCT) and stored in - 80℃ until sectioning.

### Spinal lesions

Some experiments required lesioning or recording from a slip of the ventrolateral funiculus (VLF), and other spinal segments were isolated via transection. These manipulations were performed following the removal of the spinal cord from the vertebral column, and meninges were removed prior to spinal transection, or VLF cut to prevent compression damage. The VLF was exposed by making a cut in the ventrolateral aspect of the spinal cord just caudal to the L3 ventral root. Cuts were made with ultra-fine micro-clippers (Fine Science Tools: Cat. No: 15300-00). Transected segments (thoracolumbar and sacral or thoracic and lumbosacral) were individually transported to a recording chamber where ventral roots or the VLF were outfitted with tight-fitting suction electrodes.

### Electrophysiology

Extracellular electrophysiological recordings were obtained from ventral roots, the VLF or dorsal roots with tight-fitting suction electrodes fashioned from polyethylene tubing (PE50). Signals were pre-amplified (10x) and by second stage amplifiers (Cornerstone EX4-400 Quad Differential Amplifier) at 100x for a total amplification of 1000x. Amplified signals were acquired in DC and digitized at 2.5 KHz (Digidata 1440A/1550B; Molecular Devices; Sunnyvale, CA). Data were acquired in Clampex 10.4/10.7 software (Molecular Devices) and saved on a desktop computer for offline analysis.

### Pharmacology

Episodic rhythmicity was evoked by bath application of dopamine hydrochloride (50 µM; Sigma-Aldrich). Nicotinic cholinergic transmission was blocked with mecamylamine (50 µM, Tocris) and presynaptic inhibition with the NKCC1 blocker bumetanide (50 µM, Tocris).

### Electrical stimulation of dorsal roots and episode entrainment

Episodes were entrained by applying 1 s pulse trains of electrical stimuli via a suction electrode to the dorsal root in the L5. Particular care was made to ensure that the suction electrode was not in contact with the dorsal surface of the spinal cord so that electrical stimuli would activate afferents and not ascending/descending tracts. Trains were applied at 20 Hz (pulse width: 0.4 ms), 1.5 and 1.2 or 1.0 x monosynaptic reflex threshold (TH: 7.2 ± 1.3 µA) recorded in the L5 ventral root and at an interval of 1.2 and 1.0 x episode cycle period (CP = 51.8 ± 15.4 s) measured between 25 and 30 minutes after application of 50 µM dopamine.

### In-situ hybridization and immunohistochemistry

Here, we used the Integrated Co-Detection Workflow which combines RNAscope^®^ *in-situ* hybridization with immunohistochemistry. Sections were cut at 14 μm, mounted directly onto Superfrost™ Plus slides (VWR, 483111-703), and stored at -80 ℃ until the day of experiments. The RNAscope^®^ Multiplex Fluorescent V2 kit (Cat. #323100) and RNA-Protein Co-Detection Ancillary kit (Cat. #323180) from Advanced Cell Diagnostics, Inc. (ACD) were used. The pretreatment protocol was performed according to manufacturer specifications for Fresh Frozen Tissue. Briefly, tissue was post-fixed in chilled 10% neutral buffered formalin (NBF), and then dehydrated via sequential washes of 50%, 70%, and 100% ethanol. The slides were incubated with hydrogen peroxide followed by incubation with neuronal nuclear protein (NeuN) primary antibody to label neuron somata, Clone A60 (Sigma, Cat. #MAB377, Lot #3061189, 1:500) at 4℃ overnight. The next day, the tissue was washed then fixed again in 10% NBF. Protease IV (Cat. #322381) was applied for 30 minutes. Then, we ran the RNAscope^®^ Multiplex Fluorescent v2 assay using the manufacturer’s protocol. Probes were hybridised to visualise Drd1 (ACD, Cat. #461901-C2), Drd2 (ACD, Cat. #406501), Drd3 (ACD, Cat. #447721-C3), Drd4 (ACD, Cat. #418171), Drd5 (ACD, Cat. #494411) mRNA targets. A series of 3 amplification steps were performed along with the development of fluorescent signal using Opal™ 520 (Akoya Biosciences, #FP1487001KT), and Opal™ 570 (Akoya Biosciences, #FP1488001KT). Immunohistochemistry Part B was then performed by incubation with, Donkey anti-mouse Alexa Fluor™ 488 (Life Technologies, Cat. #A21202, Lot #1562298, 1:1000) for 30 minutes at room temperature. Finally, nuclei were counterstained with DAPI prior to coverslipping with ProLong™ Glass Antifade (Invitrogen, Cat. #P36984).

### Microscopy and image analysis

Fluorescent images for RNAscope^®^ were acquired using a Leica TCS SP8 confocal microscope with 25x (0.95 NA) and 63X (1.4 NA) objectives. Fluorophores were excited with the 405 nm, 488 nm, 552 nm, and 638 nm lasers. Samples were detected with two low-dark-current Hamamatsu PMT detectors and two high-sensitivity hybrid detectors. Confocal images were acquired with a 2048 x 2048 pixels x-y resolution and 0.57 μm z-steps using the Leica Application Suite X software (RRID:SCR_013673) and exported using the Leica Image File (LIF) format. Images captured the NeuN and RNAscope^®^ probe signals (Drd1-5) as punctate dots. Offline image processing and scoring was performed in a semi-automated fashion using the spots function in IMARIS software (Bitplane AG, Zurich, Switzerland), the average NeuN+ cells and spot diameters were estimated for each slice. Automatic background subtraction was applied to enhance spot detection and a quality threshold was manually established to distinguish signal from background noise based on intensity and size criteria. Regions of interest (ROIs) were manually drawn for each lamina on each single slice. Distance transformation function was used to calculate the position and mean intensity data from NeuN+ cells and mRNA puncta in each lamina. Quantitative data from the spot detection process were exported from IMARIS for analysis with a custom-made python script. The number of cells per lamina was averaged per animal, and a group mean ± SD was calculated.

### Data analysis

Episodes of rhythmicity were analysed using custom-written Matlab scripts to detect episode properties, including cycle period, duration, amplitude, and lag time and phase between recording channels. Episode onset and offset were determined on DC signals that were detrended, band-pass filtered (0.01-1 Hz) and smoothed with a Gaussian weighted moving average. The peak amplitude of each detected episode was extracted from raw, unprocessed signals. Cycle period, duration, lag time and phase were then calculated from episode onset and offset times. Properties of the fast rhythm, including intra-episode rhythm frequency and power, were analysed using auto-wavelet spectral analysis for single ventral root recordings and cross-wavelet analysis for paired recordings using Spinal Core software. The phase of intra-episode bursting between two recordings was also extracted from cross-wavelet analyses. Procedures for these analyses have been previously published ^24,81,82^.

### Statistical analysis

Analyses of episodic rhythmicity were performed using one-way or 2-factor analyses of variance (ANOVA) where appropriate. Pharmacological or lesion-based manipulations of spinal circuits were compared to appropriate time-matched preparations using a one-way ANOVA, paired or unpaired *t*-test accordingly. Non-parametric equivalent tests were performed when normality or equal variance assumptions were not met. Post-hoc analyses (Parametric: Holm-Sidak; Non-parametric: Dunn’s) were performed when significant effects were detected from ANOVA (p < 0.05). Data are presented as mean ± standard deviation (SD). Phase analyses were conducted by performing Rayleigh statistics.

## References

1. Kjaerulff, O., and Kiehn, O. (1996). Distribution of networks generating and coordinating locomotor activity in the neonatal rat spinal cord in vitro: a lesion study. J. Neurosci. 16, 5777–5794.

2. Cowley, K.C., and Schmidt, B.J. (1997). Regional distribution of the locomotor pattern-generating network in the neonatal rat spinal cord. J. Neurophysiol. 77, 247–259.

3. Dougherty, K.J., Zagoraiou, L., Satoh, D., Rozani, I., Doobar, S., Arber, S., Jessell, T.M., and Kiehn, O. (2013). Locomotor rhythm generation linked to the output of spinal shox2 excitatory interneurons. Neuron 80, 920–933.

4. Li, E.Z., Garcia-Ramirez, D.L., and Dougherty, K.J. (2019). Flexor and Extensor Ankle Afferents Broadly Innervate Locomotor Spinal Shox2 Neurons and Induce Similar Effects in Neonatal Mice. Front. Cell. Neurosci. 13, 452.

5. Lin, S., Hari, K., Black, S., Khatmi, A., Fouad, K., Gorassini, M.A., Li, Y., Lucas-Osma, A.M., Fenrich, K.K., and Bennett, D.J. (2023). Locomotor-related propriospinal V3 neurons produce primary afferent depolarization and modulate sensory transmission to motoneurons. J. Neurophysiol. 130, 799–823.

6. Lin, S., Li, Y., Lucas-Osma, A.M., Hari, K., Stephens, M.J., Singla, R., Heckman, C.J., Zhang, Y., Fouad, K., Fenrich, K.K., et al. (2019). Locomotor-related V3 interneurons initiate and coordinate muscles spasms after spinal cord injury. J. Neurophysiol. 121, 1352–1367.

7. Chopek, J.W., Nascimento, F., Beato, M., Brownstone, R.M., and Zhang, Y. (2018). Sub-populations of Spinal V3 Interneurons Form Focal Modules of Layered Pre-motor Microcircuits. Cell Rep. 25, 146–156.e3.

8. Song, J., Ampatzis, K., Björnfors, E.R., and El Manira, A. (2016). Motor neurons control locomotor circuit function retrogradely via gap junctions. Nature 529, 399–402.

9. Bhumbra, G.S., and Beato, M. (2018). Recurrent excitation between motoneurones propagates across segments and is purely glutamatergic. PLoS Biol. 16, e2003586.

10. Lamotte d’Incamps, B., Bhumbra, G.S., Foster, J.D., Beato, M., and Ascher, P. (2017). Segregation of glutamatergic and cholinergic transmission at the mixed motoneuron Renshaw cell synapse. Sci. Rep. 7, 4037.

11. Özyurt, M.G., Nascimento, F., Brownstone, R.M., and Beato, M. (2024). On the origin of F-wave: involvement of central synaptic mechanisms. Brain 147, 406–413.

12. Nascimento, F., Özyurt, M.G., Halablab, K., Bhumbra, G.S., Caron, G., Bączyk, M., Zytnicki, D., Manuel, M., Roselli, F., Brownstone, R., et al. (2024). Spinal microcircuits go through multiphasic homeostatic compensations in a mouse model of motoneuron degeneration. bioRxiv. 10.1101/2024.04.10.588918.

13. Lambert, A.M. (2016). Dopaminergic Control of Locomotor Patterning during Development: A Tail for the Ages. Front. Cell. Neurosci. 10, 95.

14. Wiggin, T.D., Peck, J.H., and Masino, M.A. (2014). Coordination of fictive motor activity in the larval zebrafish is generated by non-segmental mechanisms. PLoS One 9, e109117.

15. Wiggin, T.D., Anderson, T.M., Eian, J., Peck, J.H., and Masino, M.A. (2012). Episodic swimming in the larval zebrafish is generated by a spatially distributed spinal network with modular functional organization. J. Neurophysiol. 108, 925–934.

16. Wahlstrom-Helgren, S., Montgomery, J.E., Vanpelt, K.T., Biltz, S.L., Peck, J.H., and Masino, M.A. (2019). Glutamate receptor subtypes differentially contribute to optogenetically activated swimming in spinally transected zebrafish larvae. J. Neurophysiol. 122, 2414–2426.

17. Montgomery, J.E., Wahlstrom-Helgren, S., Vanpelt, K.T., and Masino, M.A. (2021). Repetitive optogenetic stimulation of glutamatergic neurons: An alternative to NMDA treatment for generating locomotor activity in spinalized zebrafish larvae. Physiol Rep 9, e14774.

18. Gabriel, J.P., Ausborn, J., Ampatzis, K., Mahmood, R., Eklöf-Ljunggren, E., and El Manira, A. (2011). Principles governing recruitment of motoneurons during swimming in zebrafish. Nat. Neurosci. 14, 93.

19. Currie, S.P., and Sillar, K.T. (2018). Developmental changes in spinal neuronal properties, motor network configuration, and neuromodulation at free-swimming stages of Xenopus tadpoles. J. Neurophysiol. 119, 786–795.

20. Currie, S.P., Combes, D., Scott, N.W., Simmers, J., and Sillar, K.T. (2016). A behaviorally related developmental switch in nitrergic modulation of locomotor rhythmogenesis in larval Xenopus tadpoles. J. Neurophysiol. 115, 1446–1457.

21. Picton, L.D., and Sillar, K.T. (2016). Mechanisms underlying the endogenous dopaminergic inhibition of spinal locomotor circuit function in Xenopus tadpoles. Sci. Rep. 6, 35749.

22. Marchetti, C., and Nistri, A. (2001). Neuronal bursting induced by NK3 receptor activation in the neonatal rat spinal cord in vitro. J. Neurophysiol. 86, 2939–2950.

23. Gozal, E.A., O’Neill, B.E., Sawchuk, M.A., Zhu, H., Halder, M., Chou, C.-C., and Hochman, S. (2014). Anatomical and functional evidence for trace amines as unique modulators of locomotor function in the mammalian spinal cord. Front. Neural Circuits 8, 134.

24. Sharples, S.A., and Whelan, P.J. (2017). Modulation of Rhythmic Activity in Mammalian Spinal Networks Is Dependent on Excitability State. eNeuro 4. 10.1523/ENEURO.0368-16.2017.

25. Sharples, S.A., Burma, N.E., Borowska-Fielding, J., Kwok, C.H.T., Eaton, S.E.A., Baker, G.B., Jean-Xavier, C., Zhang, Y., Trang, T., and Whelan, P.J. (2020). A dynamic role for dopamine receptors in the control of mammalian spinal networks. Sci. Rep. 10, 16429.

26. Sharples, S.A., Parker, J., Vargas, A., Milla-Cruz, J.J., Lognon, A.P., Cheng, N., Young, L., Shonak, A., Cymbalyuk, G.S., and Whelan, P.J. (2021). Contributions of h- and Na/K Pump Currents to the Generation of Episodic and Continuous Rhythmic Activities. Front. Cell. Neurosci. 15, 715427.

27. Mahrous, A.A., and Elbasiouny, S.M. (2017). SK channel inhibition mediates the initiation and amplitude modulation of synchronized burst firing in the spinal cord. J. Neurophysiol. 118, 161–175.

28. Dalrymple, A.N., Sharples, S.A., Osachoff, N., Lognon, A.P., and Whelan, P.J. (2019). A supervised machine learning approach to characterize spinal network function. J. Neurophysiol. 121, 2001– 2012.

29. Ausborn, J., Shevtsova, N.A., Caggiano, V., Danner, S.M., and Rybak, I.A. (2019). Computational modeling of brainstem circuits controlling locomotor frequency and gait. Elife 8. 10.7554/eLife.43587.

30. Rybak, I.A., Dougherty, K.J., and Shevtsova, N.A. (2015). Organization of the Mammalian Locomotor CPG: Review of Computational Model and Circuit Architectures Based on Genetically Identified Spinal Interneurons(1,2,3). eNeuro 2. 10.1523/ENEURO.0069-15.2015.

31. Ausborn, J., Shevtsova, N.A., and Danner, S.M. (2021). Computational Modeling of Spinal Locomotor Circuitry in the Age of Molecular Genetics. Int. J. Mol. Sci. 22. 10.3390/ijms22136835.

32. Lindén, H., Petersen, P.C., Vestergaard, M., and Berg, R.W. (2022). Movement is governed by rotational neural dynamics in spinal motor networks. Nature 610, 526–531.

33. Sourioux, M., Bertrand, S.S., and Cazalets, J.-R. (2018). Cholinergic-mediated coordination of rhythmic sympathetic and motor activities in the newborn rat spinal cord. PLoS Biol. 16, e2005460.

34. Cazalets, J.-R. (2005). Metachronal propagation of motoneurone burst activation in isolated spinal cord of newborn rat. J. Physiol. 568, 583–597.

35. Falgairolle, M., and Cazalets, J.-R. (2007). Metachronal coupling between spinal neuronal networks during locomotor activity in newborn rat. J. Physiol. 580, 87–102.

36. Bonnot, A., Whelan, P.J., Mentis, G.Z., and O’Donovan, M.J. (2002). Spatiotemporal pattern of motoneuron activation in the rostral lumbar and the sacral segments during locomotor-like activity in the neonatal mouse spinal cord. J. Neurosci. 22, RC203.

37. O’Donovan, M.J., Chub, N., and Wenner, P. (1998). Mechanisms of spontaneous activity in developing spinal networks. J. Neurobiol. 37, 131–145.

38. Akay, T. (2020). Sensory Feedback Control of Locomotor Pattern Generation in Cats and Mice. Neuroscience 450, 161–167.

39. Akay, T., Tourtellotte, W.G., Arber, S., and Jessell, T.M. (2014). Degradation of mouse locomotor pattern in the absence of proprioceptive sensory feedback. Proc. Natl. Acad. Sci. U. S. A. 111, 16877–16882.

40. Bos, R., Brocard, F., and Vinay, L. (2011). Primary afferent terminals acting as excitatory interneurons contribute to spontaneous motor activities in the immature spinal cord. J. Neurosci. 31, 10184–10188.

41. Pratt, C.A., and Jordan, L.M. (1980). Recurrent inhibition of motoneurons in decerebrate cats during controlled treadmill locomotion. J. Neurophysiol. 44, 489–500.

42. Falgairolle, M., Puhl, J.G., Pujala, A., Liu, W., and O’Donovan, M.J. (2017). Motoneurons regulate the central pattern generator during drug-induced locomotor-like activity in the neonatal mouse. Elife 6. 10.7554/eLife.26622.

43. Derjean, D., Bertrand, S., Le Masson, G., Landry, M., Morisset, V., and Nagy, F. (2003). Dynamic balance of metabotropic inputs causes dorsal horn neurons to switch functional states. Nat. Neurosci. 6, 274–281.

44. Tazerart, S., Viemari, J.-C., Darbon, P., Vinay, L., and Brocard, F. (2007). Contribution of persistent sodium current to locomotor pattern generation in neonatal rats. J. Neurophysiol. 98, 613–628.

45. Masino, M.A., and Fetcho, J.R. (2005). Fictive swimming motor patterns in wild type and mutant larval zebrafish. J. Neurophysiol. 93, 3177–3188.

46. Picton, L.D., Sillar, K.T., and Zhang, H.-Y. (2018). Control of Xenopus Tadpole Locomotion via Selective Expression of Ih in Excitatory Interneurons. Curr. Biol. 28, 3911–3923.e2.

47. Cherniak, M., Etlin, A., Strauss, I., Anglister, L., and Lev-Tov, A. (2014). The sacral networks and neural pathways used to elicit lumbar motor rhythm in the rodent spinal cord. Front. Neural Circuits 8, 143.

48. Anglister, L., Cherniak, M., and Lev-Tov, A. (2017). Ascending pathways that mediate cholinergic modulation of lumbar motor activity. J. Neurochem. 142 *Suppl 2*, 82–89.

49. Bos, R., Brocard, F., and Vinay, L. (2011). Primary Afferent Terminals Acting as Excitatory Interneurons Contribute to Spontaneous Motor Activities in the Immature Spinal Cord. Journal of Neuroscience 31, 10184–10188.

50. McClellan, A.D., and Sigvardt, K.A. (1988). Features of entrainment of spinal pattern generators for locomotor activity in the lamprey spinal cord. J. Neurosci. 8, 133–145.

51. Pearson, K.G., Ramirez, J.M., and Jiang, W. (1992). Entrainment of the locomotor rhythm by group Ib afferents from ankle extensor muscles in spinal cats. Exp. Brain Res. 90, 557–566.

52. Conway, B.A., Hultborn, H., and Kiehn, O. (1987). Proprioceptive input resets central locomotor rhythm in the spinal cat. Exp. Brain Res. 68, 643–656.

53. Bracci, E., Beato, M., and Nistri, A. (1997). Afferent inputs modulate the activity of a rhythmic burst generator in the rat disinhibited spinal cord in vitro. J. Neurophysiol. 77, 3157–3167.

54. Kiehn, O., Iizuka, M., and Kudo, N. (1992). Resetting from low threshold afferents of N-methyl-D-aspartate-induced locomotor rhythm in the isolated spinal cord-hindlimb preparation from newborn rats. Neurosci. Lett. 148, 43–46.

55. Sharples, S.A., and Whelan, P.J. (2017). Modulation of Rhythmic Activity in Mammalian Spinal Networks Is Dependent on Excitability State. eNeuro 4. 10.1523/ENEURO.0368-16.2017.

56. Humphreys, J.M., and Whelan, P.J. (2012). Dopamine exerts activation-dependent modulation of spinal locomotor circuits in the neonatal mouse. J. Neurophysiol. 108, 3370–3381.

57. Bonnot, A., Chub, N., Pujala, A., and O’Donovan, M.J. (2009). Excitatory actions of ventral root stimulation during network activity generated by the disinhibited neonatal mouse spinal cord. J. Neurophysiol. 101, 2995–3011.

58. Curtis, D.R., and Ryall, R.W. (1964). NICOTINIC AND MUSCARINIC RECEPTORS OF RENSHAW CELLS. Nature 203, 652–653.

59. Curtis, D.R., Game, C.J., Lodge, D., and McCulloch, R.M. (1976). A pharmacological study of Renshaw cell inhibition. J. Physiol. 258, 227–242.

60. Mentis, G.Z., Alvarez, F.J., Bonnot, A., Richards, D.S., Gonzalez-Forero, D., Zerda, R., and O’Donovan, M.J. (2005). Noncholinergic excitatory actions of motoneurons in the neonatal mammalian spinal cord. Proc. Natl. Acad. Sci. U. S. A. 102, 7344–7349.

61. Miles, G.B., Hartley, R., Todd, A.J., and Brownstone, R.M. (2007). Spinal cholinergic interneurons regulate the excitability of motoneurons during locomotion. Proc. Natl. Acad. Sci. U. S. A. 104, 2448–2453.

62. Nascimento, F., Spindler, L.R.B., and Miles, G.B. (2019). Balanced cholinergic modulation of spinal locomotor circuits via M2 and M3 muscarinic receptors. Sci. Rep. 9, 14051.

63. Eleftheriadis, P.E., Pothakos, K., Sharples, S.A., Apostolou, P.E., Mina, M., Tetringa, E., Tsape, E., Miles, G.B., and Zagoraiou, L. (2023). Peptidergic modulation of motor neuron output via CART signaling at C bouton synapses. Proc. Natl. Acad. Sci. U. S. A. 120, e2300348120.

64. Nishimaru, H., Restrepo, C.E., Ryge, J., Yanagawa, Y., and Kiehn, O. (2005). Mammalian motor neurons corelease glutamate and acetylcholine at central synapses. Proc. Natl. Acad. Sci. U. S. A. 102, 5245–5249.

65. Calabresi, P., Picconi, B., Tozzi, A., Ghiglieri, V., and Di Filippo, M. (2014). Direct and indirect pathways of basal ganglia: a critical reappraisal. Nat. Neurosci. 17, 1022–1030.

66. Zhu, H., Clemens, S., Sawchuk, M., and Hochman, S. (2007). Expression and distribution of all dopamine receptor subtypes (D(1)-D(5)) in the mouse lumbar spinal cord: a real-time polymerase chain reaction and non-autoradiographic in situ hybridization study. Neuroscience 149, 885–897.

67. Zhu, H., Clemens, S., Sawchuk, M., and Hochman, S. (2008). Unaltered D1, D2, D4, and D5 dopamine receptor mRNA expression and distribution in the spinal cord of the D3 receptor knockout mouse. J. Comp. Physiol. A Neuroethol. Sens. Neural Behav. Physiol. 194, 957–962.

68. Briggman, K.L., and Kristan, W.B. (2006). Imaging Dedicated and Multifunctional Neural Circuits Generating Distinct Behaviors. Journal of Neuroscience 26, 10925–10933.

69. Li, W.-C., -C. Li, W., Sautois, B., Roberts, A., and Soffe, S.R. (2007). Reconfiguration of a Vertebrate Motor Network: Specific Neuron Recruitment and Context-Dependent Synaptic Plasticity. Journal of Neuroscience 27, 12267–12276.

70. Berkowitz, A. (2005). Physiology and morphology indicate that individual spinal interneurons contribute to diverse limb movements. J. Neurophysiol. 94, 4455–4470.

71. Borowska, J., Jones, C.T., Zhang, H., Blacklaws, J., Goulding, M., and Zhang, Y. (2013). Functional subpopulations of V3 interneurons in the mature mouse spinal cord. J. Neurosci. 33, 18553–18565.

72. Borowska, J., Jones, C.T., Deska-Gauthier, D., and Zhang, Y. (2015). V3 interneuron subpopulations in the mouse spinal cord undergo distinctive postnatal maturation processes. Neuroscience 295, 221– 228.

73. Zhang, Y., Narayan, S., Geiman, E., Lanuza, G.M., Velasquez, T., Shanks, B., Akay, T., Dyck, J., Pearson, K., Gosgnach, S., et al. (2008). V3 spinal neurons establish a robust and balanced locomotor rhythm during walking. Neuron 60, 84–96.

74. Danner, S.M., Zhang, H., Shevtsova, N.A., Borowska-Fielding, J., Deska-Gauthier, D., Rybak, I.A., and Zhang, Y. (2019). Spinal V3 Interneurons and Left-Right Coordination in Mammalian Locomotion. Front. Cell. Neurosci. 13, 516.

75. Chalif, J.I., Martínez-Silva, M. de L., Pagiazitis, J.G., Murray, A.J., and Mentis, G.Z. (2022). Control of mammalian locomotion by ventral spinocerebellar tract neurons. Cell 185, 328–344.e26.

76. Jean-Xavier, C., Sharples, S.A., Mayr, K.A., Lognon, A.P., and Whelan, P.J. (2018). Retracing your footsteps: developmental insights to spinal network plasticity following injury. J. Neurophysiol. 119, 521–536.

77. Sharples, S.A., and Miles, G.B. (2021). Maturation of persistent and hyperpolarization-activated inward currents shapes the differential activation of motoneuron subtypes during postnatal development. Elife 10. 10.7554/eLife.71385.

78. Sharples, S.A., Broadhead, M.J., Gray, J.A., and Miles, G.B. (2023). M-type potassium currents differentially affect activation of motoneuron subtypes and tune recruitment gain. J. Physiol. 601, 5751–5775.

79. Kyriakatos, A., Mahmood, R., Ausborn, J., Porres, C.P., Büschges, A., and El Manira, A. (2011). Initiation of Locomotion in Adult Zebrafish. J. Neurosci. 31, 8422–8431.

80. Grillner, S., McClellan, A.D., Sigvardt, K.A., Wallén, P., and Wilen, M. (1981). Activation of NMDA-receptors elicits “fictive locomotion” in lamprey spinal cord in vitro. Acta Physiol. Scand. 113, 549–551.

81. Mor, Y., and Lev-Tov, A. (2007). Analysis of rhythmic patterns produced by spinal neural networks. J. Neurophysiol. 98, 2807–2817.

82. Sharples, S.A., Humphreys, J.M., Jensen, A.M., Dhoopar, S., Delaloye, N., Clemens, S., and Whelan, P.J. (2015). Dopaminergic modulation of locomotor network activity in the neonatal mouse spinal cord. J. Neurophysiol. 113, 2500–2510.

