## Supplementary figures and images for "Episodic rhythmicity is generated by a distributed neural network in the developing mammalian spinal cord"

### Supplementary Figure 1

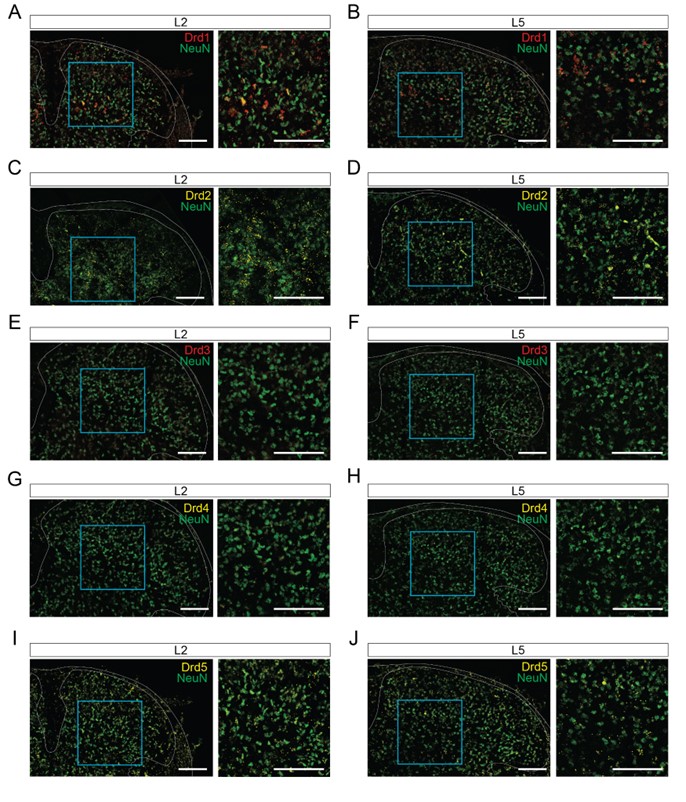

### Supplementary Figure 2

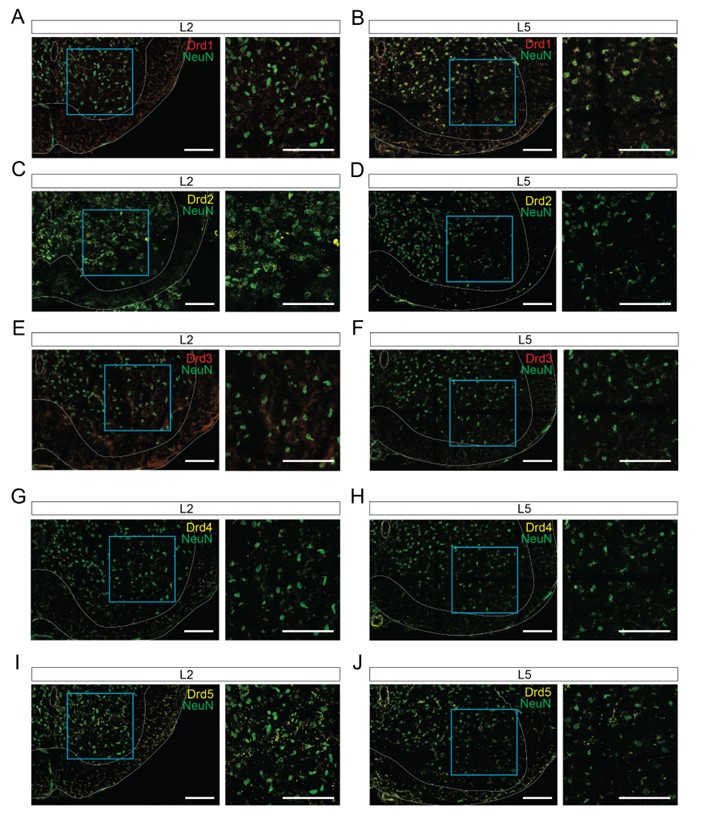

### Supplementary Figure 3

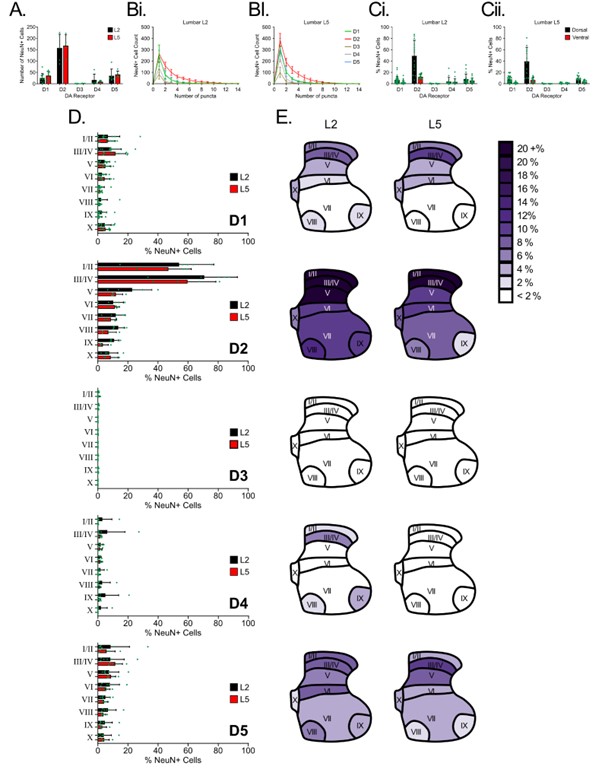
